# The transcription factor ESR2/DRNL/BOL differentially regulates *de novo* organogenesis, regeneration, and lateral root development in *Arabidopsis thaliana*

**DOI:** 10.64898/2026.09.08.744716

**Authors:** Herenia Guerrero-Largo, Beatriz Ruiz-Cortés, Ma. Isabel Cristina Elizarraraz-Anaya, Cecilia Monserrat Martínez-Pérez, Henri Manuel Jimenez-Jimenez, Joseph G. Dubrovsky, Nayelli Marsch-Martínez

## Abstract

Plant regeneration requires coordinated transcriptional and hormonal regulation to re-establish organ identity. The AP2/ERF transcription factor ENHANCER OF SHOOT REGENERATION 2 /DORNRÖSCHEN-LIKE / BOLITA (ESR2/DRNL/BOL) expressed in aerial organ founder cells promotes shoot formation, but its broader role across organogenic contexts remains unclear. Here, using loss-of-function and inducible overexpression lines of *Arabidopsis thaliana*, we demonstrate that ESR2 exerts context-dependent and antagonistic effects on shoot and root development. ESR2 activation promotes shoot and aerial-like tissue formation and enhances callus proliferation, particularly under cytokinin-rich conditions. Conversely, ESR2 suppresses or delays multiple *de novo* root formation programs, including adventitious, basal, and regenerated roots, while its loss enhances root initiation and growth. Expression analyses reveal that *ESR2* promoter activity is found at *de novo* formed basal and adventitious root primordia and emerged root apical meristems, suggesting a role in conferring regenerative competence while restricting their developmental progression. Moreover, it is expressed in newly established quiescent centers of lateral roots in intact plants and the loss of *ESR2* function strongly and negatively affects lateral root initiation, revealing an unanticipated developmental role of ESR2 and its requirement for lateral root formation. Together, these findings identify ESR2 as a shared molecular regulator governing early organogenesis in intact plants and plant explants; it establishes an aerial organ fate bias while maintaining the competence for—and limiting the progression of—*de novo* root development in explants.

## INTRODUCTION

Regeneration, defined as the dedifferentiation and re-specification process to restore or generate new structures, is a fundamental process sustained by stem cell activity. While animals display species-specific regenerative capacities, plants exhibit an extraordinary and broadly conserved ability to continuously form new organs and even regenerate entire organisms (Birnbaum and Alvarado, 2008; Heidstra & Sabatini, 2014; Liu et al., 2023).

In natural environments, this ability enables sessile plants to restore injured tissues and survive adverse environmental conditions such as herbivore attack or mechanical damage (Ikeuchi et al., 2016; Perez-Garcia & Moreno-Risueno, 2018; Sang et al., 2018). In a biotechnological context, regeneration underlies vegetative propagation methods extensively used for elite cultivars, food crops, and medicinal species (Elias et al., 2001; Gehlot et al., 2014, 2015). Many of these agronomically and medicinally important plants remain highly recalcitrant to *in vitro* regeneration, which represents a major bottleneck for their genetic improvement (Bennur et al., 2025; Luo et al., 2025). The characterization of genes and pathways regulating regeneration, or acting during regeneration, together with phytohormone signaling and wound responses, could contribute to the design of improved regeneration systems to overcome this limitation.

Phytohormones, particularly auxins and cytokinins, are central regulators of plant regeneration. The discovery of their antagonistic and synergistic roles in determining root or shoot fate laid the foundation for current plant regeneration protocols (Skoog & Miller, 1957). Their activity and distribution in the plant vary according to tissue type and developmental stage (Damodaran & Strader, 2024), and their signals act in close interaction with other regulators to determine cell identity during regeneration, such as the activity of transcription factors (Ikeuchi et al., 2017; Iwase et al., 2011, 2017).

Two types of regeneration strategies have been described: one-step (direct) and two-step (indirect). In one-step regeneration, cells undergo transdifferentiation, directly acquiring a new organ identity. In contrast, two-step regeneration involves the formation of callus tissue as an intermediate step, with organogenesis subsequently induced by hormonal cues, primarily by the ratio of auxins to cytokinins (Sang et al., 2018; Valvekens & Van Montagu, 1988).

Besides hormones (particularly auxin gradients), different studies have highlighted the roles of transcriptional reprogramming and wound signaling in this process (Bustillo-Avendaño et al., 2018; Damodaran & Strader, 2024; Ikeuchi et al., 2017). Transcriptomic and single-cell analyses have revealed dynamic transcriptional landscapes during regeneration, including the activation of pericycle-related programs (Atta et al., 2009) and cell fate transitions in leaf explants (Kareem et al., 2016; Liu et al., 2022; Sugimoto et al., 2010).

Regeneration efficiency varies depending on the plant and the explant source (Perez-Garcia & Moreno-Risueno, 2018). Roots and hypocotyls show enhanced regenerative capacity in Arabidopsis compared to other organs, due to intrinsic differences in hormonal responsiveness and cellular competence (Akama et al., 1992; Kareem et al., 2016; Valvekens & Van Montagu, 1988). In two-step regeneration protocols, the regenerative capacity also depends on the characteristics of the callus formed during the process. Calli that give rise to shoots, roots or embryos are considered organogenic and are used for plant regeneration. Conversely, calli that do not produce organs can be either friable (formed by loosely attached cells) or compact, reflecting the heterogeneous degrees of differentiation of the cells (Ikeuchi et al., 2013).

Regarding root formation, regenerated roots emerge from callus, following reprogramming through exogenous hormone application (Skoog & Miller, 1957). Furthermore, without exogenous hormones, an intact plant or plant explant can form different types of roots, depending on their origin. The primary root originates from the embryonic radicle, lateral roots arise endogenously from pericycle cells along existing roots, basal roots emerge from the hypocotyl–root junction region, denominated collet (Damodaran & Strader, 2024; Dubrovsky, 2022; Jia et al., 2019; Lucas et al., 2011; Verstraeten et al., 2014; Xu, 2018; Zobel, 2011), and adventitious roots form from non-root tissues such as hypocotyls, stems, or leaves, often in response to wounding or stress (Gonin et al., 2019; Ikeuchi et al., 2016) **(Fig. 1)**.

**Figure 1.**
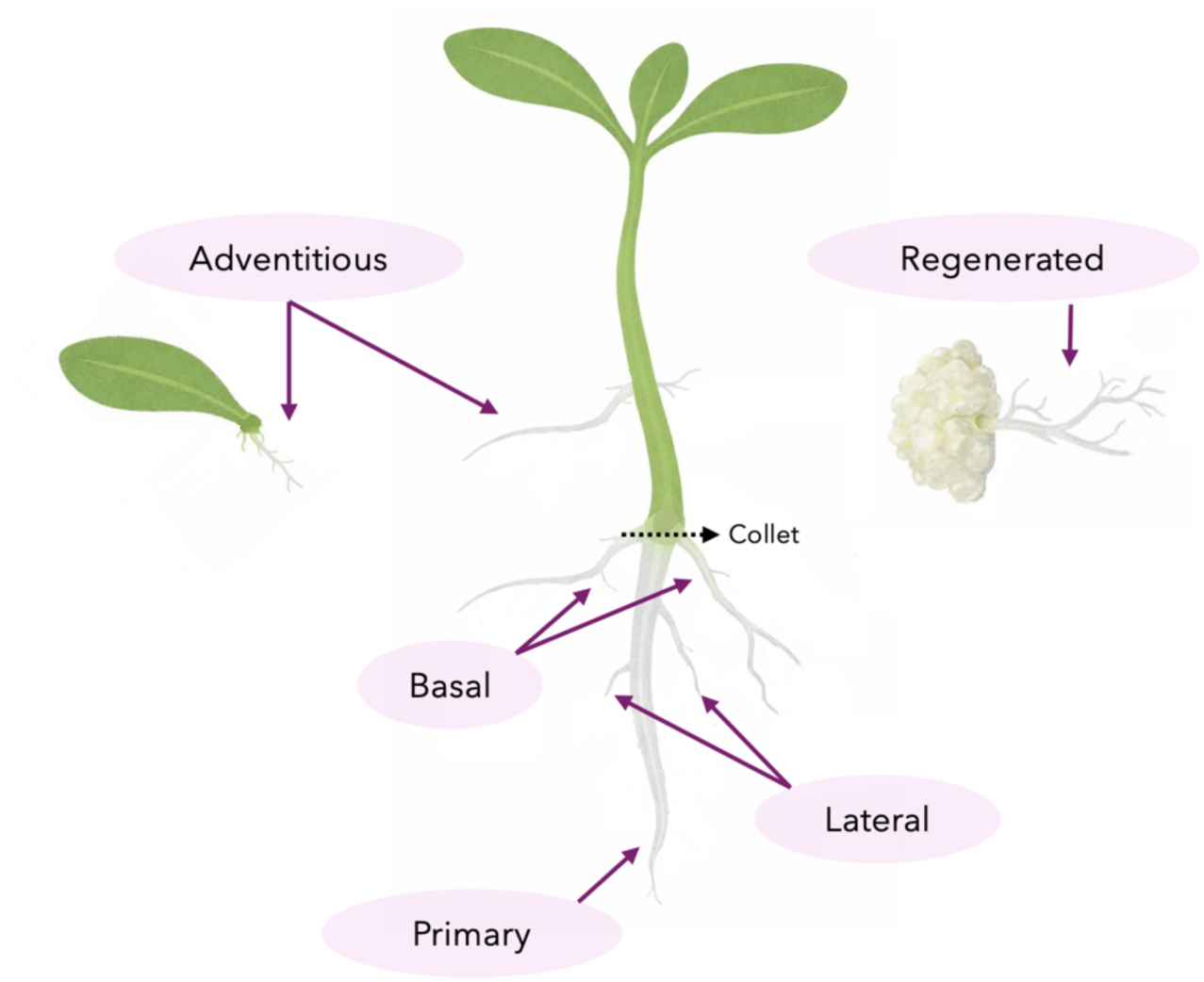
Schematic representation of distinct root types in a seedling. The primary root originates from the embryonic radicle. Lateral roots branch endogenously from the other roots. Basal roots emerge at the collet region (hypocotyl-root junction). Adventitious roots arise from non-root organs, such as the hypocotyl or leaves. Regenerated roots develops from callus tissue.

Several transcription factors, some from the APETALA2 / Ethylene Response Factor (AP2/ERF) family, have been described to participate in regeneration programs and hormonal pathways. ERF115 has been associated with wound-induced *de novo* root organogenesis from leaves (Liu et al., 2022), stem cell renewal in root meristems (Heyman et al., 2013), and indirect cytokinin pathway activation during regeneration from hypocotyl explants (Lakehal et al., 2020). *WOUND INDUCED DEDIFFERENTIATION1* (*WIND1*), another AP2/ERF member, is rapidly induced upon wounding and promotes cellular dedifferentiation in the absence of exogenous hormones (Iwase et al., 2011, 2015), acting upstream of *ENHANCER OF SHOOT REGENERATION 1 / DORNRÖSCHEN* (*ESR1/DRN*) to initiate callus formation (Iwase et al., 2017). ESR1 and ESR2, also known as DORNRÖSCHEN-LIKE and BOLITA (DRNL/BOL) (Ikeda et al., 2006; Marsch-Martínez et al., 2006; Nag et al., 2007), function in a coordinated manner during embryonic development, particularly in shoot formation (Chandler et al., 2007; Kirch et al., 2003). Their expression domains diverge during post-embryonic development: *ESR1* remains active in shoot vegetative and reproductive meristems (Kirch et al., 2003), while *ESR2* is predominantly expressed in the shoot meristem peripheral zone and is a marker of aerial organ founder cells (Chandler et al., 2011). Its expression remains in primordia and very young aerial organs, such as leaves and floral organs (Chandler et al., 2011; Marsch-Martínez et al., 2006; Nag et al., 2007). Overexpression of either *ESR1* or *ESR2* is sufficient to trigger shoot regeneration from root explants on cytokinin-rich medium (Banno et al., 2001; Ikeda et al., 2006; Matsuo et al., 2011; Nomura et al., 2009). However, ESR1 and ESR2 do not function redundantly during shoot regeneration, as the loss of *ESR2* function severely affects the shoot regeneration from root and hypocotyl explants, while in *esr1* loss-of-function mutants, shoot regeneration is barely affected (Matsuo et al., 2011).

*ESR1* has also been characterized as a WIND1 target in leaf explants (Iwase et al., 2017) that promotes auxin biosynthesis to facilitate regeneration (Lee, 2024a), while ESR2 represses auxin biosynthesis during callus induction from these explants (Lee, 2024b). ESR2 also promotes root reprogramming, causing the formation of green callus in roots (Marsch-Martínez et al., 2006) linked to the transcriptional modulation of cytokinin biosynthesis and signaling (Durán-Medina et al., 2025).

ESR2 is not only involved in shoot regeneration, but also in the formation of new organs in normal developmental contexts. Besides being a marker of founder cells in the shoot apical meristem (Chandler et al., 2011; Comelli et al., 2016), loss-of-function mutants affected in this transcription factor present malformations in organs such as cotyledons (Chandler et al., 2007) and floral organs such as stamens and gynoecia (Chandler et al., 2011; Durán-Medina et al., 2017; Nag et al., 2007; Ruiz-Cortés et al., 2025). Moreover, ESR2 and ESR1, together with REVOLUTA (REV) regulate axillary meristem initiation (C. Zhang et al., 2018). Furthermore, the loss-of-function of their closest orthologue in tomato, *LEAFLESS*, impairs leaf formation, resulting in plants that do not produce leaves (Capua & Eshed, 2017). In summary, ESR2 participates in the initiation or formation of different aerial organs.

Given the role of ESR2 in shoot regeneration and new organ formation from shoot apical meristems, this study aimed to investigate the role of ESR2 in different organogenesis contexts. Our findings suggest that ESR2 acts as a common player that regulates organogenesis differently in diverse contexts, including plant explants with and without exogenous hormone application, and intact plants. In explants, it promotes organogenetic competence favoring shoot over root identity, and at the same time delays root emergence and growth. Here we also report that in intact plants, unexpectedly, ESR2 is required for lateral root initiation.

## MATERIAL AND METHODS

### Plant materials and growth conditions

Arabidopsis *(Arabidopsis thaliana)* lines used in this study were: Wild type (WT) ecotype Columbia-0 (Col-0); transcriptional reporter line *pBOL::GUS* (Marsch-Martínez et al., 2006), mutant *bol-cr2* (Ruiz-Cortés et al., 2025), and the inducible *35S::ESR2:ER* line (Eklund et al., 2011; Ikeda et al., 2006) described in this study as *ESR2:ER*. Seeds were disinfected in a desiccator with chlorine gas (100 ml of commercial sodium hypochlorite and 5 ml of hydrochloric acid) for 4 hours. Then, the seeds were sown on 0.5x MS, 0.5% sucrose and 1.5% plant agar medium in plates. Seeds were stratified at 4°C for two days and then germinated in vertically oriented Petri dishes at 22°C under a 16 h light/ 8 h dark photoperiod. The induction of *ESR2:ER* seedlings was performed in the respective medium supplemented with 10 μM β-estradiol (β-est, Sigma-Aldrich SLBH0091V) dissolved in dimethyl sulphoxide (DMSO, Sigma-Aldrich SHBM9343, also used for mock treatments at the equivalent final concentration).

### Shoot and root regeneration from root explants

To induce shoot or root regeneration in one or two-step regeneration assays, ∼1 cm long root segments were excised from 6-day-old seedlings. The percentage of explants showing shoot/aerial-like tissue formation and root emergence, and callus area, were evaluated 20 days after explant placement in SIM or RIM.

The composition of each medium was as follows. CIM: Gamborg’s B5 salts, 2% glucose, Gamborg’s B5 vitamins, 0.5 mg/l 2,4-dichlorophenoxyacetic acid (2,4-D), 0.5 mg/l kinetin and 1.5% agar; SIM: MS salts, 1% sucrose, Gamborg’s B5 vitamins, 0.3 mg/l indole butyric acid (IBA), 0.5 mg/l trans-zeatin and 1.5% agar; RIM: MS salts, 1% sucrose, Gamborg’s B5 vitamins, 0.3 mg/l indole butyric acid (IBA) and 1.5% agar.

### Regeneration from hypocotyl explants

Regeneration from hypocotyl explants derived from 6-day-old seedlings was conducted using one-and two-step protocols. In the one-step protocol, the hypocotyl explants were directly cultured on MS medium supplemented with 3 mg/l 6-benzylaminopurine (BA). In the two-step protocol, the explants were first incubated for 8 days in MS medium containing 1 mg/l 2,4-dichloro-phenoxyacetic acid (2,4-D) and then were transferred to BA-containing medium (based on Iwase et al., 2015). Phenotypes and the percentage of explants showing shoot or root formation were evaluated 20 days after transfer to BA-containing medium.

### Leaf explant *de novo* root organogenesis assay

*De novo* root organogenesis assays were conducted using the first two rosette leaves of 12-day-old seedlings (based on Chen et al., 2014; Lee et al., 2024a; Zhang et al., 2019). Leaves were excised with intact petioles and placed with the adaxial side up onto solid B5 medium (Gamborg B5 basal medium, 0.5 g/l MES, 5% sucrose, 1.5% agar, pH 5.7). Plates were incubated under continuous light at 22°C. *De novo* root emergence was monitored at regular intervals. The proportion of leaf explants that showed root emergence, callus and the length of the longest emerged root were quantified after 15 days.

### Root formation assay using de-rooted shoot explants with and without collet

The collet region in Arabidopsis was defined in this work after (Sliwinska et al., 2012), namely as the root base region near the root-hypocotyl junction with densely located root hairs formed from each epidermal cell (a region about ∼1mm from the root base). Six-day-old seedlings were de-rooted by excising the root either with or without the collet region. Aerial explants were subsequently transferred to B5 medium and incubated under long-day light conditions. After root excision, new root formation was monitored daily in explants with and without the collet. The proportion of explants showing root emergence was evaluated 2, 3 and 6 days after excision, and the length of the longest root was measured 6 days after excision and transfer to B5 medium.

### Explant phenotypic analyses

Explant images were obtained with a Leica DM750 microscope coupled with an ICC50 HD camera and a Zeiss Stemi 2000-C stereoscope coupled with an Axiocam ERc 5s camera. Root length and callus area were measured with the ImageJ 1.53e software.

### Analysis of lateral root development

Plants grew in 0.5xMS medium supplemented with 0.5 Suc, 0.8% Bacto agar, pH 5.7. For lateral root analyses in intact seedlings, analyses were performed in plants 6 days after germination (dag) for Col and *bol-cr2* genotypes. Roots were cleared as in (Malamy & Benfey, 1997b). *ESR2:ER* seeds were sown in 0.5x MS medium, and 4 dag seedlings were transferred to the same medium supplemented with 10 µM β-estradiol. They were analyzed before and after the transfer.

### Analysis of *ESR2* expression using the *pBOL::GUS* marker line

For GUS staining, seedlings or explants were collected in 1 ml of 90% cold acetone and were fixed under vacuum for 5 minutes. Then, they were incubated for 2 hours in 90% acetone at room temperature. Acetone was replaced with 1 ml X-gluc solution (100 mM Sodium Phosphate Buffer pH7, 10 mM EDTA pH 8, 5 mM Potassium Ferricyanide, 5 mM Potassium Ferrocyanide, 1 mg/ml X-gluc diluted in DMSO) and vacuum was applied for 5 minutes. They were afterwards incubated in X-gluc solution at 37°C overnight.

Seedlings or explants were cleared and hydrated with a series of ethanol (70%, 40%, 20%, 10%) for 10 minutes each at room temperature, ending with a 1 ml 50% glycerol 2% DMSO solution, and were mounted on slides for microscopy. Whole-mount preparations were analyzed with a Leica DMR microscope equipped with Nomarski differential interference contrast (DIC) optics and a Leica DFC420C digital camera or with Olympus BX53 microscope equipped with DIC optics and CoolSNAPcf CCD Photometrics Camera.

### *ESR2* expression analyses in publicly available databases

The GEO dataset GSE161970, corresponding to the study “Transcriptional dissection of lateral root primordium cells” in *Arabidopsis thaliana* (Serrano-Ron et al., 2021) was used to explore *ESR2/BOL/DRNL* expression during lateral root formation. The raw count matrix was imported into R (version 4.4.2, R Core Team, 2021) and subsequently converted into an analysis object using the Seurat package. The data were normalized using the standard NormalizeData method. Cell identities were assigned based on annotations previously included in the dataset (column V25), which correspond to relevant cell types in the root. For this analysis the pericycle, and lateral root primordia (LRP) were chosen. The expression of *ESR2* was evaluated and compared with *WOX5*, as a root quiescent center marker (Sarkar et al., 2007; Tian et al., 2014). Normalized expression values per cell were extracted, and expression averages were calculated by cell type. To facilitate visualization, conditional formatting was applied in the form of a heatmap, where the values were represented on a continuous color scale (white to blue). Finally, the data were analyzed using log-normalized values, not raw counts, where 0 corresponds to “not detected”, 0.01–0.1 to “very low”, 0.1–0.5 to “low/moderate” and >1 to “high” expression. The *Arabidopsis thaliana* Tiling Array Express (Laubinger et al., 2008) and the Root Cell Atlas (Root Cell Atlas Initiative, Department of Plant Systems Physiology, 2022) were also included in the analysis.

### Statistical analysis

Statistical analysis and graphs of all data in the study were conducted through R (version 4.4.2, R Core Team, 2021), Rstudio (Version 4.4.2, Posit team, 2025), or SigmaPlot, version 12 (Systat Software, Grafiti LLC, Palo Alto, CA, USA). Each statistical test used and the number of repetitions are indicated in the legends of the corresponding figure. For dichotomous data, test for binomial proportions was used. For parametric variables, Student’s *t* test was used.

## RESULTS

To investigate the role of ESR2/DRNL/BOL in organogenesis, we tested different explant sources in a variety of conditions and intact seedlings. For consistency, the names of the genotypes used are based on their originally published designation. We analyzed wild type Col-0 (WT), loss-of-function *(bol-cr2,* Ruiz-Cortés et al., 2025) and inducible activation lines (*ESR2:ER* in the text, where ESR2 entrance to the nucleus is induced by β-estradiol; Ikeda et al., 2006; Eklund et al., 2011).

### *ESR2* promotes cell reprogramming and inhibits root formation in root explants

First, root explant regeneration was tested. One-step and two-step regeneration assays were performed employing either shoot inducing medium (SIM) or root inducing medium (RIM) as the only or the last step **(Fig. 2a and b)**. For the one-step assay, root explants were directly placed onto shoot-inducing (SIM) or root-inducing (RIM) media (based on the protocol described by Ikeda et al., 2006). For the two-step regeneration assay, root explants were first cultured on callus-inducing medium (CIM) to promote cellular dedifferentiation and subsequently transferred to SIM or RIM (based on Iwase et al., 2017).

**Figure 2.**
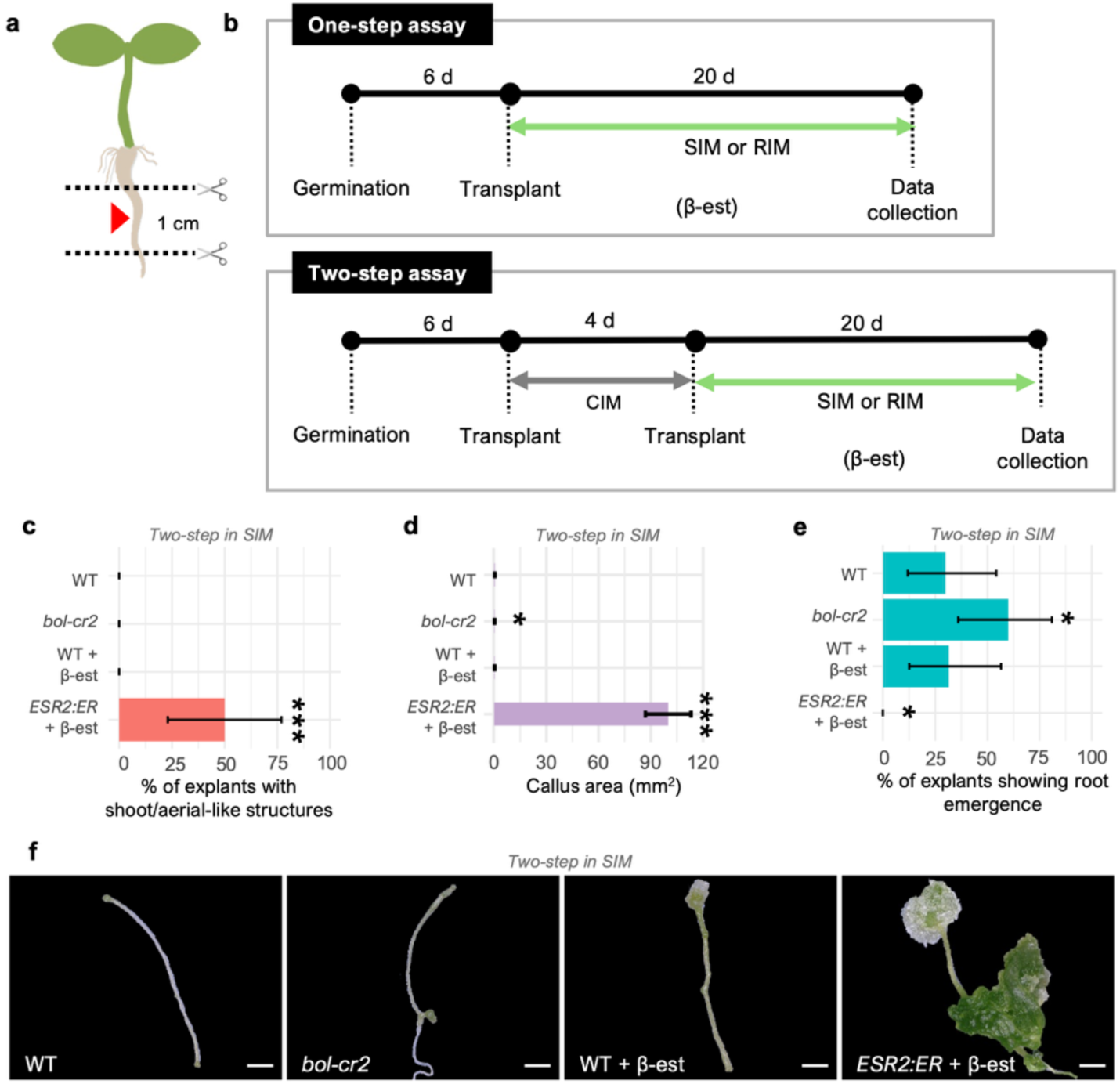
Effects of ESR2 in one and two-step regeneration assays using root explants. **a** Schematic of the region from where root explants (∼1 cm) were obtained, used for regeneration assays in Shoot Inducing Medium (SIM) or Root Inducing Medium (RIM) with one-step and two-step protocols. The red arrowhead indicates the excised root segment. **b** Schematic representation of the timeline and treatments applied in one-step (top) and two-step (bottom) SIM or RIM regeneration assays. *ESR2:ER* was induced with β-estradiol (β-est) during the SIM or RIM transplant phase. Data collection was performed at the end of the 20-day SIM or RIM incubation. **c–f** Results from the two-step assay after transfer to SIM. **c** Frequency of explants showing shoot/aerial-like regeneration. Error bars indicate 95% confidence intervals calculated using an exact binomial method (* *p*<0.05, *** *p*<0.001, test for binomial proportions). **d** Callus area. Bars represent the mean. Error bars represent SE (* *p*<0.05, *** *p<*0.001, Student’s *t-*test). **e** Frequency of explants showing root emergence in the two-step assay. Error bars indicate 95% confidence intervals calculated using an exact binomial method (* *p*<0.05, test for binomial proportions). f Representative root explants of each genotype. c-f n = 16–20 seedlings per line, two independent experiments. Scale bars = 1 mm.

The rate of shoot regeneration from root explants transferred directly to SIM was analyzed first. In the one-step assay, shoot regeneration was not detected in WT, WT+β-est and *bol-cr2* **(Fig. S1a)**. Interestingly, the *ESR2:ER* line showed shoot regeneration, as well as differentiated tissues that did not correspond to shoots *per se*, but rather resembled other types of aerial tissue denominated in this study as “aerial-like” regeneration that includes leaf, stem and inflorescence-like formation. After *ESR2* induction, 36.4% of the explants presented shoot or aerial-like regeneration **(Fig. S1a and d)**. Callus formation occurred across all genotypes, but *ESR2:ER* explants formed significantly larger calli (∼12 mm²) with distinctive green and sometimes vitrified morphology, in contrast to the smaller (0.4-1 mm²), whitish/yellowish calli observed in other lines **(Fig. S1b and d)**. Finally, the percentage of *bol-cr2* and *ESR2:ER* explants that showed root emergence ranged between 9 and 28%, with no significant differences compared to WT explants **(Fig. S1c)**.

In the two-step assay, the pattern of regeneration of aerial-like “organs” from root explants transferred to SIM was consistent with the one-step assay: only *ESR2* induction triggered the formation of shoot/aerial-like structures with a frequency of 50% **(Fig. 2c)**. Additionally, callus development was also dramatically increased in *ESR2:ER* line after induction, with callus area approaching 100 mm^2^ **(Fig. 2d)**, and these calli exhibited aerial-like structures **(Fig. 2f)**. Unexpectedly, the loss of *ESR2* function significantly enhanced the proportion of root explants that showed the emergence of at least one root (60%; **Fig. 2e)** compared to WT (30%). In contrast, ESR2 induction strongly diminished the proportion of root explants that presented root emergence (0%) compared with WT+β-est explants (32%).

In summary, *ESR2* overexpression promoted the development of shoot/aerial-like tissues and led to a marked increase in callus proliferation, particularly in the two-step regeneration assay. Conversely, the absence of *ESR2* function enhanced root organogenesis in the two-step regeneration assay, highlighting its antagonistic role in root and shoot regeneration. These results suggest that ESR2 plays a significant role in promoting aerial identity organogenesis in all cases tested in this assay, while suppressing root formation in particular conditions.

Next, we analyzed root explants of WT, *ESR2* loss-of-function mutant *(bol-cr2)* and the inducible line *(ESR2:ER),* again using the one-and two-step regeneration assays, but now transferring the explants to the auxin-rich Root Inducing Medium (RIM) conditions. The assay followed the same timing and treatment conditions as for SIM-based assays **(Fig. 2b)**, allowing for comparison in distinct hormonal contexts.

In both one-step and two-step assays with root explants, ESR2 induction led to a reduction in the percentage of explants showing root organogenesis compared to controls **(Figs. S2a and S3a)**, indicating a consistent inhibitory effect on root organogenesis even in auxin-rich conditions. Notably, ESR2 induction also triggered the formation of shoot-like or aerial-like structures in a small subset of explants; their frequency remained low but statistically significant compared to the WT **(Fig. S2b and S3b)**.

Callus proliferation was markedly enhanced in explants where ESR2 activity was induced **(Fig. S2c and S3c**), reaching areas up to ∼12 mm² in the one-step and ∼18 mm² in the two-step assay. The resulting calli displayed prominent green coloration and occasionally exhibited disorganized outgrowths resembling partially differentiated aerial tissues **(Fig. S2d and S3d)**. In contrast, other genotypes showed limited callus expansion (0.4-0.6 mm² in one-step and 0.03-1.5 mm² in two-step assays) with no evident organogenetic activity.

Finally, the *bol-cr2* mutant did not exhibit major differences compared to the WT in any measured parameter, suggesting that loss of *ESR2* alone is insufficient to disrupt auxin-mediated regeneration programs. These results indicate that the loss of *ESR2* function does not significantly affect regeneration capacity in auxin-rich conditions. However, ESR2 increased activity antagonized the auxin-induced root emergence program, suppressing root organogenesis and promoting enhanced cell proliferation at callus state (leaning towards aerial identities, given the green color of calli).

### *ESR2* induction promotes the formation of structures with partial shoot identity and hampers root formation in hypocotyl explants

We next investigated the role of ESR2 during regeneration using WT, loss-of-function *(bol-cr2)* and inducible *(ESR2:ER)* hypocotyl explants **(Fig. 3a)**. Both one-and two-step cytokinin-based assays were performed **(Fig. 3b)**.

**Figure 3.**
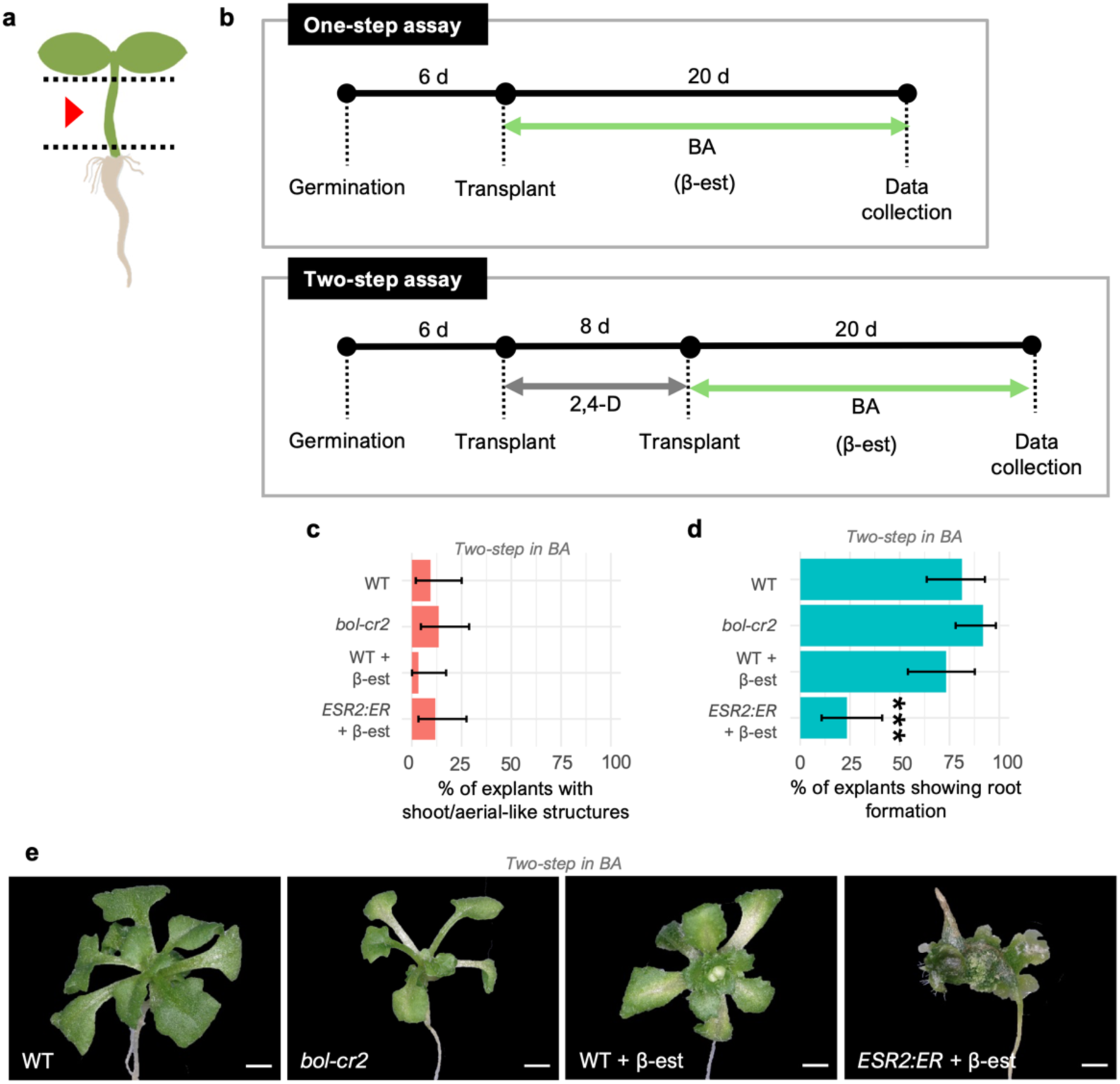
Effects of ESR2 in regeneration assays from hypocotyl explants. **a** Schematic of hypocotyl explants used for Benzylaminopurine (BA) regeneration assays in one-step and two-step protocols. The red arrowhead indicates the excised hypocotyl segment. **b** Schematic representation of the BA regeneration assays. *ESR2:ER* was induced with β-estradiol (β-est) during the BA transplant phase. Data collection was performed at the end of the 20-day BA incubation. **c–e** Results from the two-step assay after transfer to BA. c-d Frequency of explants showing shoot/aerial-like (c) and root formation (d). Error bars indicate 95% confidence intervals calculated using an exact binomial method (*** *p*<0.001, test for binomial proportions). N = 32-37 seedlings per line, two independent experiments. **e** Representative phenotypes of shoot/aerial-like regeneration in each genotype. Scale bars = 1 mm.

In the one-step protocol, all genotypes showed comparable percentages of explants presenting callus or some kind of tissue or organ formation **(Fig. S4a and S4b)**. However, *ESR2:ER* explants formed prominent green calli with irregular morphology upon induction, suggesting early reprogramming events without complete shoot development **(Fig. S4c)**.

In the two-step assay, *bol-cr2* hypocotyl explants did not differ from WT in either shoot or root formation frequencies **(Fig. 3c and d)**. However, the percentage of explants showing root emergence was markedly reduced in induced *ESR2:ER* explants **(Fig. 3c)**. Despite similar frequencies of shoot/aerial-like regeneration across genotypes, the identity and morphology of the regenerated tissues varied. WT explants produced well-formed shoots, with some serrated leaves, while in *bol-cr2* explants leaf size was noticeably smaller **(Fig. 3e)**, although regeneration was visible at similar time points across genotypes. In *ESR2:ER* induced explants, the regenerated structures were often disorganized and included green calli tissues, which were counted as aerial-like organs to calculate the frequency of regeneration.

In summary, ESR2 induction promotes the formation of proliferative green tissues and disorganized shoot/aerial-like tissues, suggesting an alteration in regenerative properties. Rather than producing fully differentiated shoots, ESR2-induced hypocotyl explants developed compact green calli and abnormal aerial structures, consistent with an intermediate regenerative state between proliferating callus and organized shoot formation. Moreover, ESR2 induction suppressed root regeneration in hypocotyl explants under the conditions studied. Conversely, *ESR2* loss-of-function did not significantly alter regenerative outcome frequency in this context, only impacting the regenerated shoot organ size.

### ESR2 inhibits *de novo* root formation in leaf explants

Next, we studied *de novo* root formation from WT, *bol-cr2*, and *ESR2:ER* leaf explants **(Fig. 4a)**. This assay does not include any hormonal treatment (Chen et al., 2014; Lee et al., 2024a; G. Zhang et al., 2019). Explants were transferred to Gamborg B5 medium directly after excision, and data were collected 15 days post-transfer **(Fig. 4b)**.

**Figure 4.**
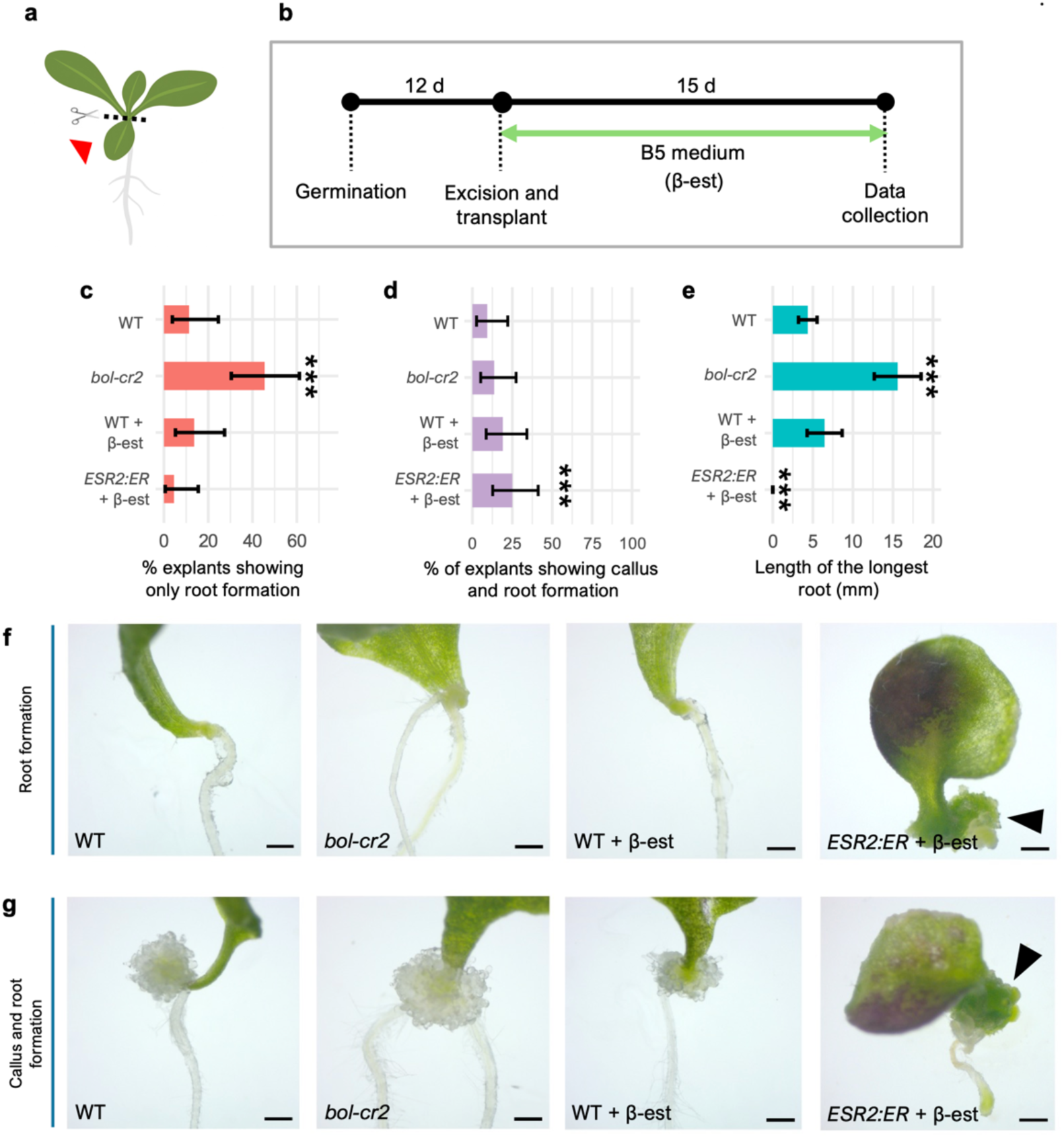
Effects of ESR2 in regeneration from leaf explants. **a** Schematic of the leaf explants used. The red arrowhead indicates the excised leaf. **b** Schematic representation of the regeneration assay. *ESR2:ER* was induced with β-estradiol (β-est) in the Gamborg B-5 (B5) medium phase. Data collection was performed at the end of the 15-day B5 incubation. **c-d** Proportion of explants showing adventitious root formation (c) and callus and root formation (d). Error bars indicate 95% confidence intervals calculated using an exact binomial method (*** *p*<0.001, test for binomial proportions). **e** Length of the longest adventitious root. Error bars indicate SE (*** *p*<0.001, Student’s *t*-test). **f-g** Representative images of explants of each genotype showing adventitious root (f) and callus and root (g) formation after 6 days after excision (dae). *ESR2:ER* explants showed callus, and when roots formed, they exhibited green sectors. Black arrows indicate green calli formation. c-e n = 44 seedlings per line, two independent experiments. Scale bars = 1 mm.

In these assays, *de novo* roots were formed in all genotypes, although the frequency of explants that showed adventitious roots, and the morphology and characteristics of the organs or tissues formed varied. In some explants, only roots were observed **(Fig. 4c)**, while others showed both roots and calli **(Fig. 4d).** Approximately 42% of *bol-cr2* leaf explants developed roots directly, compared to 10% in WT and ∼5% in *ESR2:ER* explants induced with β-est **(Fig. 4c).** ∼13% *bol-cr2* leaf explants displayed roots associated with callus, comparable to WT **(Fig. 4d).** By contrast, ∼25% of the induced *ESR2:ER* explants showed a higher frequency of roots associated with green calli, statistically different to control explants, in which the formed calli were also different (white and smaller; **Fig. 4d).** Root length measurements revealed that *bol-cr2* explants developed significantly longer roots (∼15 mm) than WT (∼5 mm), whereas *ESR2:ER* explants induced displayed severely stunted roots (0 mm; **Fig. 4e**). In some cases, *ESR2:ER* explants formed small green calli with no visible root outgrowth, which could be due to a similar reprogramming effect of ESR2 overexpression in root tissues (**Fig. 4f and g**; Durán-Medina et al., 2025). Notably, explants that developed roots from the petiole seldom simultaneously produce green callus, and *vice versa*, indicating mutually exclusive outcomes.

In summary, the loss of *ESR2* enhances both *de novo* root formation frequency and root length, while its overexpression strongly suppresses root development, indicating that ESR2 functions as a negative regulator of adventitious root formation in leaf explants.

### *ESR2* is expressed prior to *de novo* root emergence in leaf explants

Since *ESR2* caused a reduction in root formation from leaf explants, we monitored its expression after leaf excision and during adventitious root formation using the transcriptional reporter line *pBOL::GUS* (Marsch-Martínez et al., 2006). Leaf explants were obtained and processed as for the previous experiment (as described in **Fig. 4a and b)**, and GUS staining was performed at 1 to 5, 10 and 15 days after excision (dae) and transfer to B5 Gamborg medium.

No GUS signal was observed 24 h after excision **(Fig. 5a and b)**, but was first detected at 2 dae **(Fig. 5c)**. At this time 66.6% of the analyzed explants displayed two small expression domains adjacent to the vasculature, located slightly above the cut site, 13.3% presented an expression domain expanded towards the wound site, coinciding with regions of visibly enlarged cells **(Fig. 5d, inset)**, and the remaining explants (∼20%) showed no detectable expression at this stage. At 3 dae, 25% of the explants exhibited bidirectional expansion of the expression domain along the vasculature **(Fig. 5e)**, whereas in most explants (75%) expression was primarily restricted to the wound-proximal region and elongated cells **(Fig. 5f, inset).** A similar pattern was observed at 4 dae, 30% of explants showed staining expanding both upwards the proximal end of the cut site (**Fig. 5g**), while 70% maintained expression mainly near the cut region **(Fig. 5h)**.

**Figure 5.**
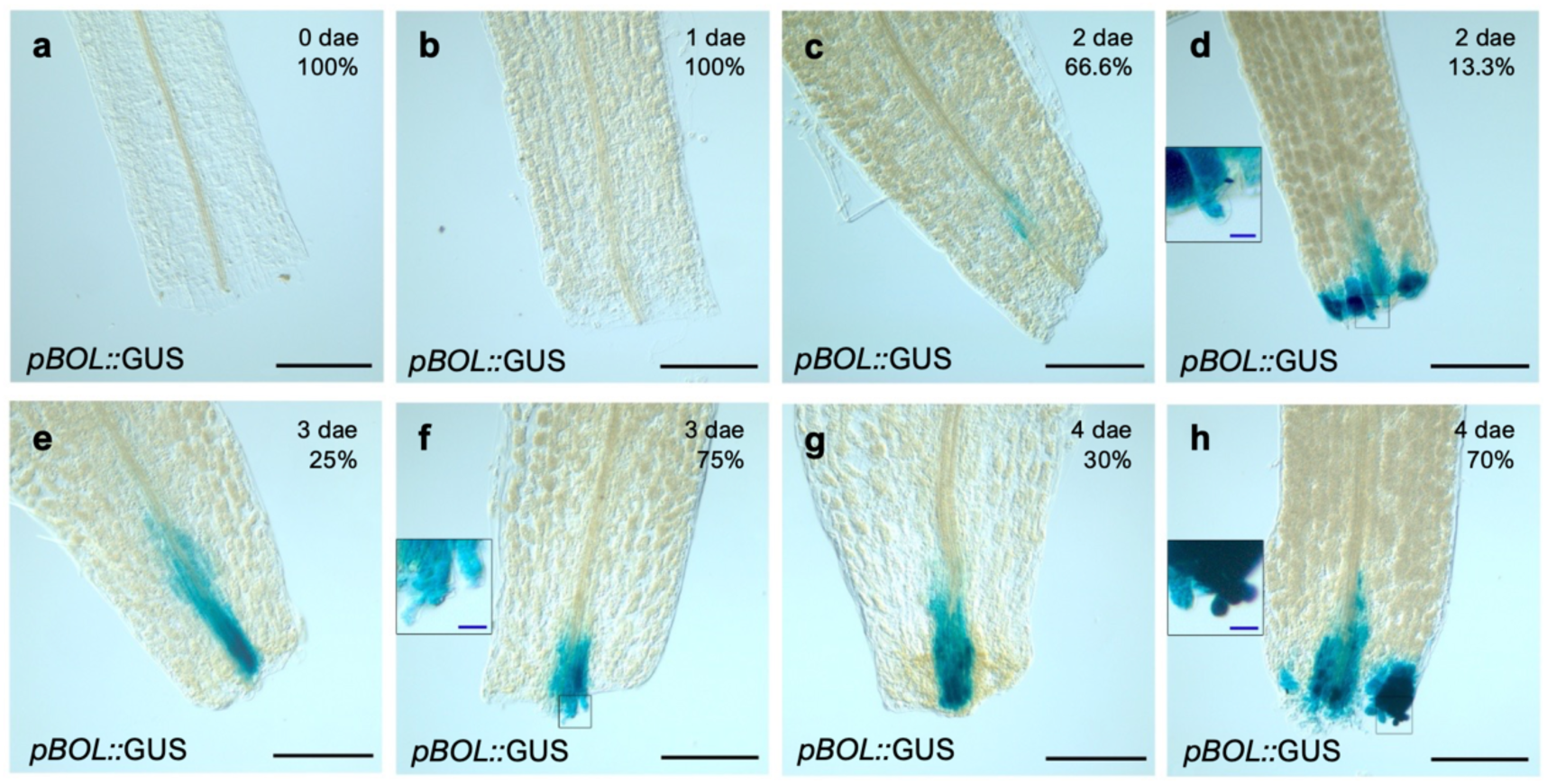
*ESR2* expression during adventitious root formation from leaf explants. *ESR2* expression evaluated in the *pBOL::GUS* reporter line at **a**, zero; **b**, one; **c-d**, two; **e-f**, three and **g-h**, four days after excision (dae) and transference onto B5 Gamborg’s medium. Insets on d, f and h show a magnified view of the boxed region, highlighting elongated cells. The percentage out of a total of 15 explants that presented each staining pattern is shown below the days. Scale bars: black = 0.2 mm; blue = 20 µm.

Five dae, GUS signal was still observed at the vasculature region just above the cut petiole stump, ending as a localized point at the center of the stump edge **(Fig. S5a).** Ten dae, approximately 72% of the explants exhibited an expansion of the expression domain, at the edge of the cut petiole. This region could represent a transitional tissue, emerging callus, or new adventitious root primordia **(Fig. S5b).** At fifteen days, all explants (100%) displayed strong GUS staining **(Fig. S5c and d),** with enriched staining in the proximal region of the leaf explant, at what seemed to be the region of a callus closest to the leaf, and the tips of adventitious roots.

These results reveal that *ESR2* is not directly activated as an early wound response gene, but progressively during the organogenesis process, with expression at intermediate and late stages of root formation, when cell proliferation and identity transitions become more evident. The spatial pattern and the effect of the loss of its function suggest a role in regenerative domains and potentially restricting adventitious root development.

### *ESR2* delays basal root emergence and adventitious root growth from aerial explants while providing competence for root initiation in the hypocotyl

After observing that loss of *ESR2* function increased the proportion of explants that formed adventitious roots, while overexpression reduced it, we evaluated its function in root organogenesis from de-rooted young aerial explants **(Fig. 6a).** We tested different seedling ages and transfers to different media after root excision, as summarized in **Table 1**. Moreover, de-rooted explants with or without collet (a hypocotyl -root junction as defined in Materials and Methods, **Fig 1**) were included in the comparison. As little is known about nature of basal roots (also called “anchor roots”; Jia et al., 2019) we addressed basal root emergence from the explants with collet region and the role of ESR2 in their development **(Table 1**, **Fig. 6a).** The evaluated genotypes included again the loss-of-function mutant *(bol-cr2)* and its control (WT), the ESR2 inducible overexpression line (*ESR2:ER*+β-est) and its corresponding controls (*ESR2:ER*, WT+Mock, *ESR2:ER*+Mock, and WT+β-est). We evaluated the percentage of plants that formed basal and/or adventitious roots and measured the length of the longest root.

**Figure 6.**
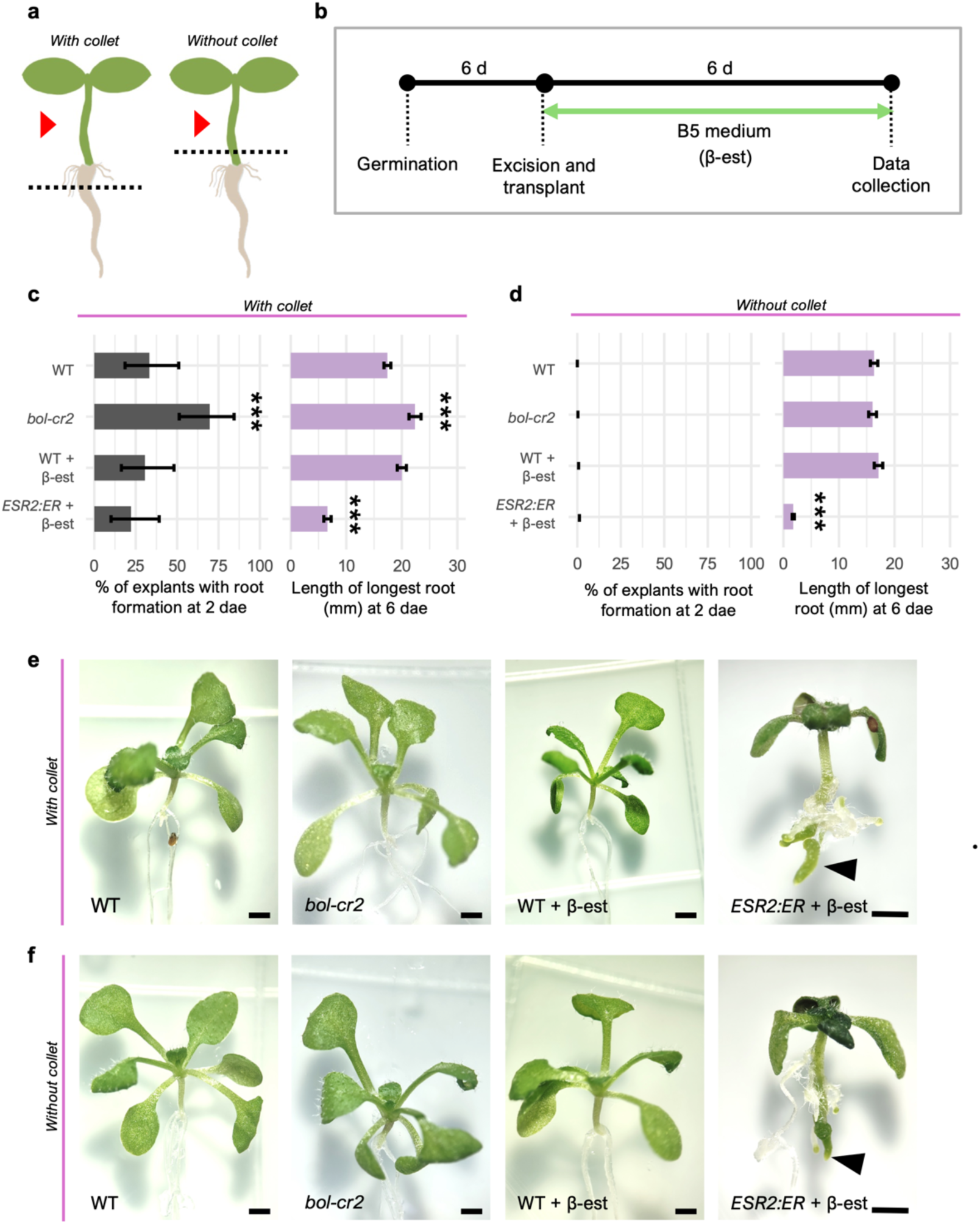
Effects of ESR2 in root organogenesis from de-rooted shoot explants. **a** Schematics of the de-rooted aerial explants used. The red arrowhead indicates the excised shoot segment with (left) and without (right) collet. **b** Schematic representation of the regeneration assay. *ESR2:ER* was induced with β-estradiol (β-est) during the Gamborg B-5 (B5) medium phase. Data collection was performed daily after transplanting. **c-d** Comparison of the frequency of root formation 2 days after excision (dae), and length of the longest root 6 dae with (c) and without (d) collet. For root formation frequency, error bars indicate 95% confidence intervals calculated using an exact binomial method (*** *p*<0.001, test for binomial proportions). For root length, error bars indicate SE (*** *p*<0.001, Student’s *t*-test). n = 33-36 seedlings per line, two independent experiments. **e-f** Representative explant images of different genotypes with (e) and without (f) collet at 6 dae. Black arrows indicate green calli formation. Scale bars = 1 mm.

**Table 1.** Experimental conditions evaluated for the standardization of the *de novo* root formation assay from aerial explants. β-est: β-estradiol; dag: days after germination. B5: Gamborg B-5.

| Genotype/Treatment | Explants | Plant age from the explant | Culture Medium | Evaluated Variables |
| --- | --- | --- | --- | --- |
| WT<br>*bol-cr2*<br>*ESR2:ER*<sup>a</sup><br>WT+Mock<sup>a</sup><br>*ESR2:ER*+Mock<sup>a</sup><br>WT+β-est<br>*ESR2:ER*+β-est | With collet<br>Without collet<br>Only collet | 6 dag<br>9 dag<sup>b</sup> | MS<sup>b</sup><br>B5 | Frequency of root formation<br>Length of longest root |

<sup>a</sup> No statistically significant differences were observed between these lines and the WT+β-est (β-estradiol) control.
<sup>b</sup> Conditions/treatments with inconsistent results across independent replicates.

In the initial tests for protocol standardization, we analyzed basal and adventitious root formation in shoot explants obtained from seedlings aged 6 and 9 days after germination (dag). In explants derived from 9-dag seedlings the timing and frequency of root development varied considerably, so only 6-day-old seedlings were finally evaluated **(Fig. 6b)**. We also compared MS and Gamborǵs B5 media, and the latter was chosen for the analysis because differences among genotypes were more evident **(Fig. 6b).** The initial explants analyzed included aerial tissue retaining the collet, aerial tissue with the collet removed, and the collet alone. In some cases, calli and *de novo* root formation were observed in the collet alone; however, growth eventually ceased, suggesting a loss of viability or progressive explant death. Consequently, subsequent analyses focused only on aerial explants with or without the collet **(Fig. 6a).** Under our chosen experimental conditions, we observed that when the collet was included in the explant, basal roots began to emerge as early as two days post-excision **(Fig. 6c).** This was faster than the previous assays using other tissues or hormones, that required from 10 to up to 20 days to observe roots. The *bol-cr2* mutant exhibited a significantly higher frequency of explants exhibiting root development compared to the WT control **(Fig. 6c, left panel).** Conversely, the ESR2 inducible line did not show significant differences compared to its control (WT+β-est). Six days after excision (dae), all explants had roots. Accordingly, the *bol-cr2* loss-of-function mutant developed substantially longer roots compared to its control, whereas the overexpression line displayed markedly shorter roots **(Fig 6c, right panel).** At this evaluated time, therefore, ESR2 delays root formation and growth in aerial explants that include the collet.

We also analyzed the type and number of roots formed per explant with collet six dae. Curiously, at this time *bol-cr2* and *ESR2:ER* showed a higher total number of roots per explant compared with their controls, but the number of basal (emerged from the collet) and adventitious (emerged from the hypocotyl) roots per explant largely differed between genotypes **(Fig. S6a and b)**. All genotypes formed basal roots **(Fig. 6e)** and *bol-cr2* explants displayed a higher number of basal roots than WT explants **(Fig. S6a)**. In contrast, WT and *bol-cr2* explants with collet did not form any adventitious root at the evaluated time. Curiously, from all genotypes, only induced *ESR2:ER* explants developed adventitious roots **(Fig. S6a and b)**.

Next, we analyzed shoot explants without collet. Interestingly, in contrast with those explants with collet, root formation was not observed during the first two dae and transplantation **(Fig. 6d**, left panel). New roots became evident only until the third day **(Fig. S6c).** In this context, the *bol-cr2* mutant did not show significant differences in either adventitious root formation frequency three dae **(Fig. S6c)** or root length compared to WT explants six dae **(Fig. 6d, right panel).** However, the induced *ESR2:ER* line showed reduced frequency of root formation three dae **(Fig. S6c)** and shorter roots relative to their control six dae **(Fig. 6d, right panel).** At this time, all explants had visible roots, and curiously, the position at which roots emerged along the hypocotyl in these explants lacking the collet also differed in different genotypes.Adventitious roots only formed close to the cut surface in all genotypes except in the *ESR2:ER* induced line, where roots were distributed along the entire hypocotyl **(Fig. 6f).**

In both explants, with or without collet, green calli were occasionally observed at the base of the hypocotyl or the new roots **(Fig. 6e and f).** This indicates that ESR2 also enhances cell proliferation and apparently causes partial reprogramming in the developing new root primordia, similar to what is observed in the roots of intact plants (Durán-Medina et al., 2025).

In summary, when comparing the data obtained from explants with and without collet, the collet appears to inhibit the formation of adventitious roots in wild type and *bol-cr2* explant a few days after primary root excision. Increased ESR2 activity, conversely, promotes adventitious root formation even in the presence of the collet. Without the collet, explants show adventitious root formation only at the base of the hypocotyl. However, ESR2 also promotes adventitious root formation at upper regions of the hypocotyl. Moreover, ESR2 appears to naturally delay basal root development, and its induction delays adventitious root growth the first dae, but roots develop afterwards.

### *ESR2* is expressed prior to *de novo* basal and hypocotyl-derived adventitious root growth

To investigate the *ESR2* spatio-temporal expression pattern in shoot explants during new root organogenesis, we analyzed the *pBOL::GUS* reporter line comparing seedlings before cutting, and aerial explants with and without collet at one, two, and three days after primary root excision (dae). *GUS* expression became detectable in the collet region of intact seedlings around three days after germination (dag) in approximately 70% of seedlings, increased to 80% by four dag, and was consistently observed in all seedlings by six dag **(Fig. 7a to e).** Notably, expression was detected in the core domain of basal root primordia with the strongest signal corresponding to the prospective quiescent center **(Fig. 7c to e)**.

**Figure 7.**
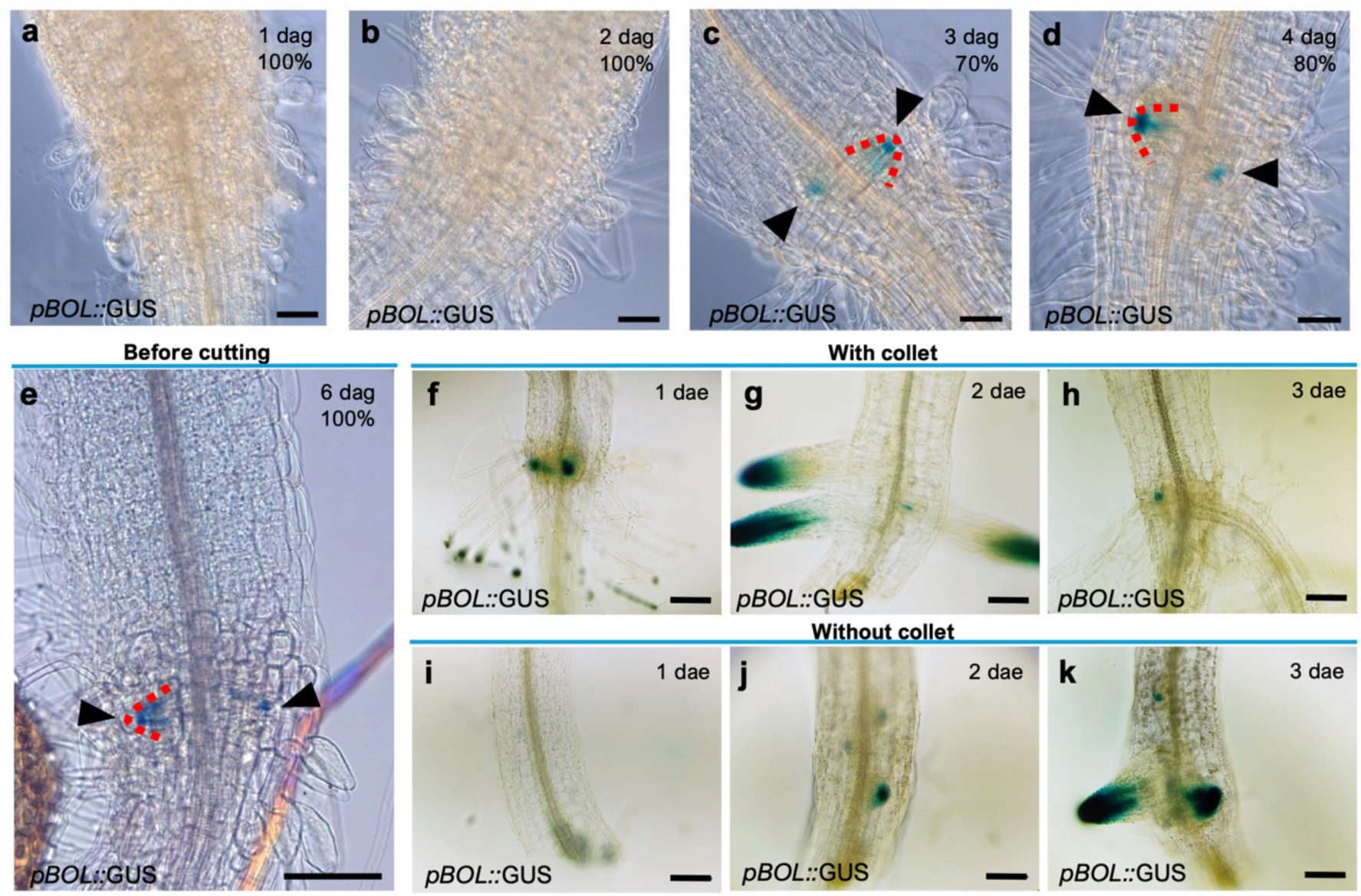
*ESR2* expression in seedlings and aerial explants with or without collet. **a-d** *ESR2* expression in the collet region of intact plants 1–4 days after germination (dag). The percentage out of a total of 10 seedlings presenting expression is indicated under the number of dag. **e** *ESR2* expression in the collet region prior to excision, 6 dag. **f-h** *ESR2* expression in de-rooted explants with collet region 1–3 days after excision (dae). **i-k** *ESR2* expression in aerial explants lacking the collet region 1–3 dae. Black arrowhead = GUS signal. Red dashed line delimits basal root primordia. Scale bars a-d = 50 μm, e-k = 100 μm.

In cut explants with collet, after excision *ESR2* expression was maintained in the primordia of basal roots and then was localized in the root apical meristem of developing basal roots **(Fig. 7f).** Two and three dae, expression was sustained at the tip of emerged basal roots and new root primordia **(Fig. 7g and h).**

In explants lacking collets, *ESR2* expression was absent in the hypocotyl of aerial explants one dae **(Fig. 7i).** Interestingly, two dae, *ESR2* expression became detectable at the inner regions of the hypocotyl above the excision zone, in the tips of developing <u>adventitious</u> root primordia **(Fig. 7j).** By three dae, *ESR2* expression increased and was observed at the tips of emerging adventitious roots and as discrete points at the inner tissues of the hypocotyl above the new roots, possibly corresponding to new root primordia **(Fig. 7k).**

In summary, *ESR2* expression is consistently localized at the tips of basal and adventitious root primordia and in the apical meristems of newly formed basal and adventitious roots.

### ESR2 regulates early lateral root development in intact plants

The differences in basal and adventitious root formation after root excision opened the question about the role of *ESR2* in lateral root (LR) formation, and this was addressed using the loss-of-function mutant *bol-cr2* and the overexpressor *ESR2:ER* line. To this end, we estimated LR primordium (LRP) densities within the LR formation zone, that comprises the longitudinal region of the primary root between the first detected LRP, closest to the root tip and the first emerged LR (Dubrovsky & Forde, 2012) in intact plants of the different genotypes. The LRP density in the *bol-cr2* mutant was significantly lower than that in WT roots **(Fig. 8a).** Similarly, the density of all LR initiation events (including LRPs and LRs) was also significantly decreased in the mutant line **(Fig. 8b).** Because differences in LR density in some cases can be explained by differences in the length of primary root cells, the LR initiation index should be estimated (Dubrovsky et al., 2009). For this, we measured the length of 10 cortical cells per root across five roots per genotype and no significant differences were detected (**Fig. 8c,** p > 0.05, Student’s *t*-test), indicating no changes in LR initiation index, therefore confirming reduced LR initiation in *bol-cr2*. In addition to the effect of the mutation on LR initiation, we also estimated how the mutation affected LR emergence. While in 6 dag WT plants 45% of all initiation events resulted in LR emergence, in the mutant it was only 23 % (*n* = 11-16, *p* < 0.001, Student’s *t*-test). This analysis suggests that LRP morphogenesis is slower in the mutant compared to WT. In two out of 11 six dag mutant plants LRs were not found at all, supporting this conclusion. Overall, this analysis suggests that *ESR2* is required for LR initiation and morphogenesis.

**Figure 8.**
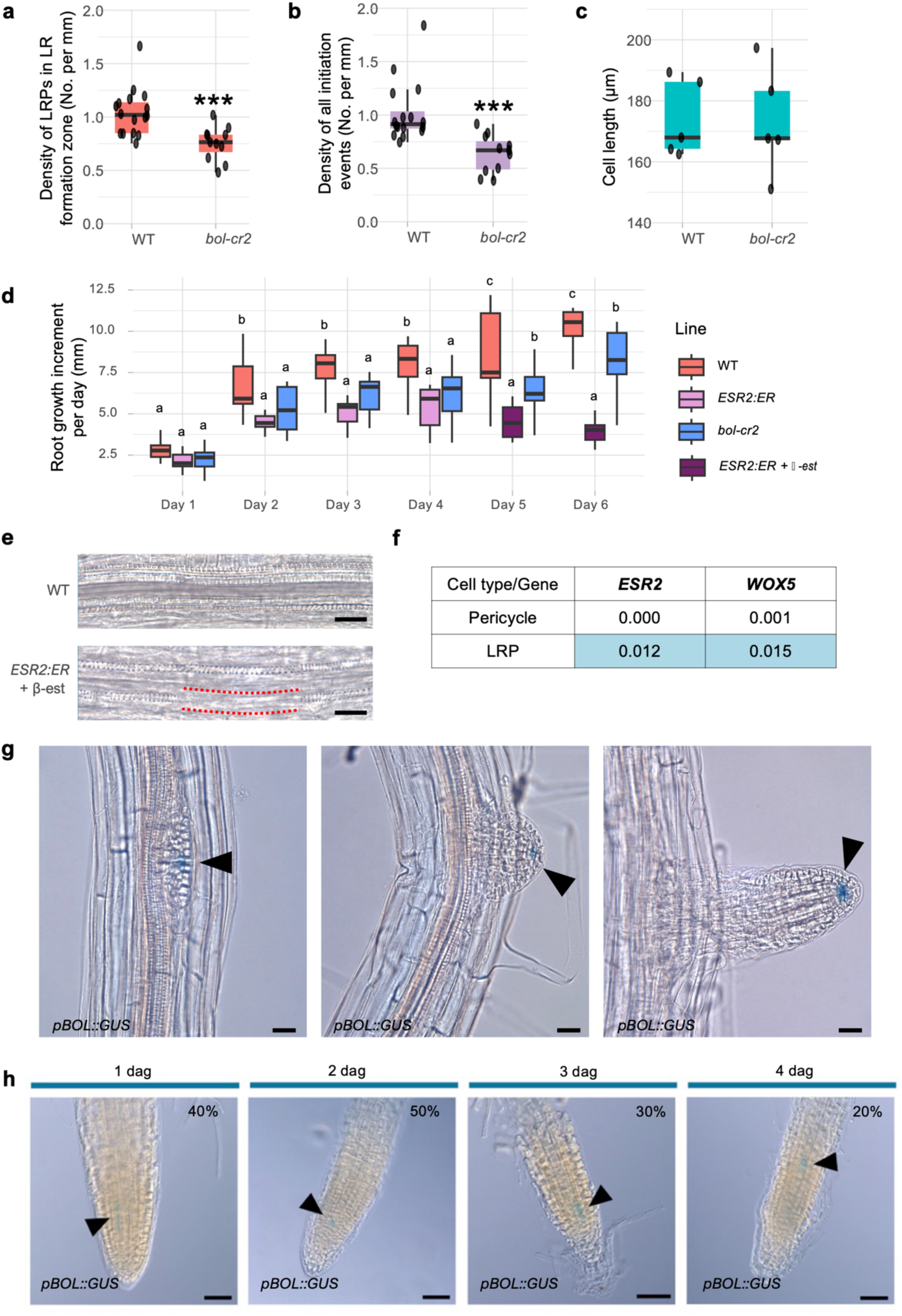
*ESR2* role in lateral root development in intact seedlings and its expression pattern. **a** Density of lateral root primordia (LRPs) within the lateral root (LR) formation zone in WT and *bol-cr2* plants (n = 11– 16, *p*<0.001, Mann–Whitney Rank Sum Test). **b** Density of total lateral root initiation events (LRI, including LRPs and emerged lateral roots) in WT and *bol-cr2* (n = 11–16, *p*<0.001, Mann–Whitney Rank Sum Test). **c** Length of fully elongated cortical cells in WT and *bol-cr2* (n = 10 cells per root, 5 roots per genotype, two-tailed *p*=0.941, Student’s *t*-test). **d** Daily root growth increment in WT, *ESR2:ER*, and *bol-cr2* seedlings during six days of observation. ESR2:ER activity in the *ESR2:ER* line was induced by β-estradiol (β-est) at 5 and 6 days. Letters indicate significant differences among lines within each day according to Tukey’s HSD test (*p*<0.05). **e** Longitudinal views of cleared roots showing protoxylem structure in WT and lack of protoxylem in induced *ESR2:ER* line. Red dashed lines indicate protoxylem discontinuities. Scale bars = 20 μm. **f** *ESR2* expression level in different root cell types and comparison with the quiescent center (QC) marker *WOX5.* Data obtained from GSE161970 public dataset (Serrano-Ron et al., 2021), values represent mean log-normalized expression levels obtained using the Seurat NormalizeData function (scale factor = 10,000). Values are expressed in arbitrary units (a.u.). **g** *ESR2* expression in lateral root primordia and emerged lateral roots at different developmental stages, indicated by black arrowheads. Scale bars = 20 μm. **h** *ESR2* expression in the primary root tip 1–4 days after germination (dag), indicated by black arrowheads. Percentages indicate the proportion of seedlings showing detectable GUS signal (n = 10). Scale bars = 50 μm.

We next examined the effect of *ESR2* overexpression on LR initiation. After two days of β-estradiol (β-est) induction of 4 dag *ESR2:ER* plants, LR initiation was strongly inhibited within the root portion formed in presence of β-est. Among seventeen roots analyzed in two independent experiments, five showed no LRP initiation, four produced only one LRP, seven produced two LRP, and one produced three LRP. Curiously, quantification of initiation events in the root portion formed prior to induction (4 dag plants) revealed a significantly lower density compared to WT controls (it was 0.4 ± 0.3 LR initiation events per mm in the *ESR2:ER* line whereas in WT this density at comparable time period was 1.0 ± 0.1 events per mm, mean ± SD, *p* < 0.001, Student’s *t*-test; *n* = 16-17), suggesting some activity leakage, as previously observed (Durán-Medina et al., 2025). Root growth increased over time in all three genotypes, although the magnitude of the increase differed among lines **(Fig. 8d).** During the first dag, no significant differences were detected among WT, *ESR2:ER*, and *bol-cr2* plants. From day 2 onwards, WT generally exhibited the highest growth values, whereas *ESR2:ER* roots displayed the lowest growth rates. By days 5 and 6 (after *ESR2:ER* induction), all three genotypes differed significantly from each other, with WT showing the highest growth rates and *ESR2:ER* the lowest. This growth inhibition in *ESR2:ER* roots was accompanied with abnormalities in protoxylem development; in five out of seventeen roots within the root portion formed in presence of β-est, interrupted protoxylem was found **(Fig. 8e).** Therefore, changes in ESR2 levels affect correct LRP formation, and its overexpression inhibits parent root growth and development.

Given the LR phenotypes in the loss-of-function mutant, we explored publicly available single cell transcriptome datasets of developing LRP (Serrano-Ron et al., 2021) to find out whether *ESR2* was expressed during early stages. Indeed, reads corresponding to *ESR2* were found in cells labelled as LRP **(Fig. 8f)** at an equivalent level to *WOX5*, expressed at the quiescent center of both in primary (Sarkar et al., 2007), and lateral roots (Tian et al., 2014). The expression level of both genes was close to the detection threshold, suggesting expression is restricted to specific cellular subpopulations during LRP development, as known for *WOX5*. Moreover, very low *ESR2* expression in roots was also found in the *Arabidopsis thaliana* Tiling Array Express (Laubinger et al., 2008) and the Root Cell Atlas (Root Cell Atlas Initiative, 2022). Interestingly, analysis of *ESR2* promoter activity, using the *pBOL::GUS* reporter line showed that it is strictly limited to the QC of developing LRs. Promoter activity was found starting from StIV LRPs, when the QC is known to be specified (Goh et al., 2016) and in the QC of recently emerged growing LRs **(Fig. 8g)**.

To find out whether *ESR2* was also expressed at the QC of root apical meristems of primary roots, its promoter activity was evaluated in developing roots of seedlings 0 to 4 days after germination. Interestingly, while no expression at the QC was detected, it was observed at a distance of 3 to 6 cortical cells shootward of the QC in the internal root apical meristem tissues, putatively in the pericycle, and it was maintained only within a short region corresponding to 2 to 6 cortical cells **(Fig. 8h)**. The proportion of seedlings displaying this expression pattern decreased over time, being detected in approximately 40% of seedlings at day 0, 50% at day 1, 30% at day 2, and 20% at day 4.

In summary, *ESR2* is expressed during LRP formation and in young LRs. Its loss-of-function or a higher expression level inhibited LR initiation and delayed LR emergence. Overall, the data supports a previously unknown role for *ESR2* in LR initiation and morphogenesis.

## DISCUSSION

Organogenesis, the process of coordinating patterning, morphogenesis and differentiation of cells into functional organs, is crucial in multicellular organisms that have organs performing different functions (Andrews & Priya, 2025). Intact plants are capable of reiteratively forming numerous postembryonic organs (Perianez-Rodriguez et al., 2014). New organs can also form after wounding or the application of exogenous hormones, a process known as regeneration, that can even give rise to a new plant from a plant explant (Birnbaum & Alvarado, 2008). There are common genetic factors that participate in new organ formation both in intact plants, and in wound-induced or regeneration processes, such the WUS/WOX transcription factors and auxin activities among others (Adem et al., 2024; Ikeuchi et al., 2022; Rasheed et al., 2024). In intact Arabidopsis plants, the ESR2/BOL transcription factor marks founder cells that give rise to new organs in the shoot apical meristem and is also required for hormone-promoted shoot regeneration from explants (Chandler et al., 2011; Matsuo et al., 2011). In this study, ESR2 was found to have contrasting functions and effects in hormone-induced regeneration and wound-triggered organogenesis in an organ and context-dependent manner **(Fig. S7).** Moreover, an unexpected role for ESR2 in lateral root development in intact plants and basal root development in shoot explants with collet was identified.

### ESR2 overrides hormone-induced regeneration programs and modulates organogenesis in a context-dependent manner

The regeneration of a whole plant from a plant fragment requires the reprogramming of its cells to form new meristems and organs (Ikeuchi et al., 2016, 2019; Kareem et al., 2016; Sugimoto et al., 2010). The hormones auxin and cytokinin aid in this process, and their ratio promotes the induction of callus that can direct towards root (high auxin) or shoot (high cytokinin) formation (Skoog & Miller, 1957). Under cytokinin-rich conditions, *ESR2* overexpression induced the formation of disorganized calli and shoot-like structures, suggesting a shift in cellular identity toward an aerial fate. This is consistent with previous reports indicating that high cytokinin-to-auxin ratios promote shoot identity, and that *ESR* genes can enhance shoot regeneration when ectopically expressed in root-derived tissues (Banno et al., 2001; Ikeda et al., 2006). Shoot regeneration is a continuum of developmental states (Shin et al., 2020) that includes the establishment of a functional SAM, stable WUS-CLV signaling, and subsequent leaf initiation and outgrowth (Gordon et al., 2007). Partial regeneration represents incomplete reprogramming of tissues, in which shoot identity markers are activated but boundary formation, correct patterning, hormonal gradients, or stem cell niche stabilization are insufficient (Gordon et al., 2007). *ESR2* expression promoted shoot identity but appeared to limit meristem maturation. In our regeneration assays, *ESR2* overexpression promoted shoot-like regeneration under cytokinin-rich conditions but frequently resulted in incomplete or abnormal aerial structures. This suggests that ESR2 may guide shoot identity acquisition without fully supporting later phases required for sustained meristem maintenance and organization, and leaf development. These results align with previous reports showing that *ESR* genes enhance shoot induction efficiency but can disrupt proper meristem organization when ectopically expressed (Banno et al., 2001; Ikeda et al., 2006; Matsuo et al., 2011). Accordingly, ESR2 has been classified as a shoot pro-meristem gene (Shin et al., 2020). Given that *ESR2* is expressed at the earliest stages of aerial organ development and during regeneration, ectopic ESR2 activity could lock tissues into an intermediate developmental state, promoting shoot identity while simultaneously inhibiting later differentiation events required for organ formation. Alternatively, ESR2 activity alone may not be enough to allow progression, and additional regulators or signals may be required. High cytokinin:auxin ratios lead towards shoot and away from root identities (Pernisová et al., 2009; Skoog & Miller, 1957; Werner et al., 2003). ESR2 promotes cytokinin accumulation, while also activating the expression of the cytokinin signaling inhibitor AHP6. Notably, ESR2 increased activity in *ahp6* mutants allowed the formation of true shoots (Durán-Medina et al., 2025), pointing at an imbalance in hormonal homeostasis as a possible cause of the incomplete progression of shoot formation.

Even in auxin-prevailing conditions, typically associated with root fate determination, *ESR2* overexpression promoted green callus formation, markedly suppressing root emergence **(Figs. S2a and S3a).** For example, upon *ESR2* induction in the one-step essay, root explants did not show any root formation (**Fig. S2a**). As the LR initiation event density in *ESR2:ER* line before β-estradiol induction was 0.4 events mm^-1^, on average 2.3 LRPs are expected to be present per mm of the root explant. Complete absence of emerged LRs thus suggests that high ESR2 activity may alter the developmental trajectory of primordia, redirecting them from root toward shoot identity and arresting their progression. Future experiments combining primordium markers, clearing techniques, and time-resolved imaging can help elucidate this.

Conversely, loss of *ESR2* function led to higher rates of root formation in root explants transplanted to Shoot Inducing Medium (SIM) in the two-step assays, compared to wild type plants **(Fig. 2e and f)**. ESR2 has been reported to negatively regulate auxin biosynthesis, guiding HISTONE DEACETYLASE 6 to *YUCCA* promoters in leaf explants placed in Callus Inducing Medium (CIM, Lee et al., 2024b). Interestingly, ESR2 also positively regulates auxin biosynthesis indirectly through STYLISH1 in developing aerial organs (Eklund et al., 2011). This contrasting regulation occurs in different developmental contexts, so ESR2 appears to produce opposite effects in auxin biosynthesis in different tissues and conditions. In this case, the inhibiting role of ESR2 in root formation from hormone-treated root explants may be explained by its negative regulation of auxin biosynthesis and can be further tested.

### ESR2 modulates root organogenesis from leaf and de-rooted shoot explants

The formation of new organs is also triggered by wounding (Birnbaum & Alvarado, 2008). Arabidopsis leaf explants can form adventitious roots (Chen et al., 2014; de Klerk et al., 1999) and have aided the study of wound-induced *de novo* root organogenesis. The use of leaf explants and de-rooted shoot explants with or without the collet **(Figs. 4 to 6),** revealed a function of ESR2 in wound induced adventitious and basal root formation. Since in these explants root formation relies predominantly on endogenous hormonal signaling and wound-induced pathways, ESR2 may modulate intrinsic reprogramming pathways that guide *de novo* organogenesis and activation.

ESR2 acts as a repressor of adventitious root formation in hormone-free leaf explants, in agreement with previous reports of a different *esr2* mutant allele (Lee et al., 2024a). ESR2 is not wound-activated. Instead, ESR2 expression increased over time in the vascular region near the cut site of the leaf explants. Strong expression at earlier times in a similar tissue was reported in leaf and root explants placed in 2,4-D or CIM supplemented medium (Lee et al., 2024b). Therefore, ESR2 may act as an indicator or competence factor for new organ formation and a cell reprogramming marker that at the same time negatively affects root identity or root development progression in contexts like hormone-induced regeneration and exogenous-hormone-free leaf explants.

ESR2 could function in a two-phase manner during root regeneration from leaf explants. In the early phase, ESR2 expression may promote acquisition of regenerative competence by facilitating cell de-or redifferentiation and proliferation. This is consistent with previous reports showing that *ESR* genes are induced during early stages of organogenetic reprogramming triggered by exogenous hormones (Ikeda et al., 2006; Matsuo et al., 2011). However, *ESR2* expression may maintain reprogrammed cells in a transitional state, inhibiting or lacking other factors required for progression toward organized root formation and subsequent outgrowth.

The aerial explant approach allowed us to further uncover the role of ESR2 in the formation of basal roots and wound induced hypocotyl-derived adventitious roots **(Fig 6 and 7).** Root formation from aerial explants proved to be a sensitive and fast platform that can aid future studies investigating genetic or chemical factors involved in basal and adventitious root formation and emergence, especially in conditions where hormonal autonomy is critical. The root-hypocotyl junction, also referred to as the collet, is where basal roots are formed and emerge. *ESR2* is already expressed before wounding in basal root primordia at the collet of young seedlings. Our results suggest that ESR2 functions as a negative regulator of basal root primordium wound-induced activation within the collet. In intact seedlings, ESR2 expression in this region may maintain basal root primordia in a dormant state, as loss of *ESR2* function accelerates basal root emergence when the collet is present in de-rooted shoot explants. Upon excision of the primary root, these primordia are rapidly activated similar to the reported data after the RAM excision (Jia et al., 2019), enabling swift responses in transcriptional programs, that most likely depend on signaling perceived at the collet. When the collet was present, no adventitious roots in the hypocotyl were observed neither in *bol-cr* nor in wild type explants at the evaluated times **(Fig. 6e, S6a and b)**. Curiously, while *ESR2* overexpression reduced the proportion of aerial explants with visible roots at early times, it unexpectedly led to the formation of adventitious roots in the hypocotyls at later times **(Fig. 6e S6a and b),** suggesting again that the ectopic expression of ESR2 enables or promotes competence, in this case to form new primordia at the inner tissues of the hypocotyl. In these explants, the hypocotyl may be less sensitive to the root-progression repressive action of *ESR2*, so new root primordia were able to develop into organs similar to adventitious roots. Given the importance of an apocarotenoid, anchorene, for basal root formation (Jia et al., 2019) and its role in modulation of auxin homeostasis (Ke et al., 2024), it could be important to investigate how ESR2 effects on basal root formation are related to carotenoid metabolism in the collet region.

Removal of the collet in aerial explants eliminated both *ESR2* expression and rapid primordium activation, thereby delaying root formation until new organogenic-competent zones were established in the hypocotyl. According to a possible organogenetic competence-providing role, *ESR2* expression reappeared at later time points in collet-less explants as discrete points related to new adventitious root primordia presumably in the prospective QC region of developing primordia and then was maintained in emerging adventitious root tips (**Fig. 7i to k**).

In WT de-rooted explants without collet, adventitious root formation occurred predominantly near the excision (Damodaran & Strader, 2024). Interestingly, in the *ESR2* overexpression line, this localized pattern was disrupted, and adventitious roots emerged at upper regions along the hypocotyl (**Fig. 6f**). Because not all regions of the hypocotyl are equally competent for regeneration (Damodaran & Strader, 2024), the appearance of roots distant to the cut site again suggests that ESR2 promotes organogenesis competence at these sites. Previous studies have shown that cytokinin signaling restricts the zone of competence during adventitious root development, thereby defining the spatial limits for organogenesis (Damodaran & Strader, 2024). Notably, ESR2 has been found to dually interact with the cytokinin pathway during fruit development (Durán-Medina et al., 2017). It also promotes cytokinin biosynthesis while simultaneously promoting signaling inhibition, and both actions are required for green callus induction by ESR2 (Durán-Medina et al., 2025; Ikeda et al., 2006). Moreover, it fosters cytokinin inactivation during regeneration at the shoot apical meristem (Casimiro et al., 2001; Ikeda et al., 2006; Malamy & Benfey, 1997a; Parizot et al., 2008). Therefore, it is plausible that ESR2 acts through cytokinin signaling inhibition or inactivation to promote competence in the hypocotyl inner tissues.

Adventitious root formation competence also involves IBA-derived IAA (Damodaran & Strader, 2024). This highlights the potential for future studies to investigate the interaction between ESR2 and the cytokinin and auxin pathways during derooting-induced hypocotyl-derived adventitious root formation. Future studies could explore how ESR2 integrates hormonal cues to modulate basal and adventitious root primordium formation and *de novo* root organogenesis from aerial tissues.

### ESR2 is expressed in lateral root primordia and young lateral roots, and is required for lateral root development in intact seedlings

While adventitious roots derive from non-root tissues (Verstraeten et al., 2014) and the primary root arises from embryonic tissues, lateral roots (LRs) in Arabidopsis form postembryonically from founder cells specified in pericycle, a root inner tissue located at the periphery of the vascular cylinder (Casimiro et al., 2001; Malamy & Benfey, 1997a; Parizot et al., 2008). A close homologue of ESR2, PUCHI, is required for LR morphogenesis (Hirota et al., 2007), and *ESR2* gene dosage is important for lateral root (LR) development in intact plants, since a lack or an excess of ESR2 perturbed correct LR initiation (this work, **Fig. 8a and b**). Moreover, ESR2 expression was constrained to a few cells during middle stages of LR development. It was consistently restricted to the QC and started to be present in StIV LRPs, a stage when the QC markers are first detected during primordium formation (Goh et al., 2016), and it was maintained in emerged young lateral roots. *ESR2* is not constantly or broadly expressed at the primary root, which explains why its expression may have been overlooked, though a report of lateral root perturbation in a different *ESR2* loss-of-function mutant allele further confirms ESR2 function in LR development (Lee et al., 2024b).

The fact that *ESR2* promoter activity was found in the QC and tips of adventitious, basal, and lateral roots, suggests that it is important for LRP morphogenesis, and further studies should address this possibility. Absence of *ESR2* in the loss-of-function mutant *bol-cr2* can explain slow LRP morphogenesis. Together, this indicates that ESR2, unexpectedly, participates in lateral root formation, opening the questions of whether it plays a role in the founder cell specification during LR initiation and how exactly it affects LRP morphogenesis when expressed in the QC.

Another issue worth examining is whether the role of ESR2 in lateral root development could be related to its capacity to promote callus formation and its role in shoot regeneration (Durán-Medina et al., 2025; Ikeda et al., 2006; Marsch-Martínez et al., 2006; Matsuo et al., 2011). Hormone-induced calli have characteristics associated with LRPs, and histological events observed during shoot development resemble early LR formation (Atta et al., 2009; Cary et al., 2002; Che et al., 2002, 2007; Kareem et al., 2016; Sugimoto et al., 2010, 2019), for example, found root QC markers in the subepidermal layer of hormone-induced callus, and *ESR2* is expressed at the QC of LRP and young LRs (**Fig. 8g**). Interestingly, some known genes required for proper LR development, such as *IAA14* (Fukaki et al., 2002), *LBD16* (Goh et al., 2019) and *LRP1* (Smith & Fedoroff, 1995) are differentially expressed in root tissue after *ESR2* induction (Lazcano-Ramírez et al., 2021). Whether callus induced by ESR2 overexpression is equivalent to hormone-induced callus remains as a quest for future research.

Finally, *ESR2* is expressed at founder cells of aerial organs and remains in leaf and floral organ primordia at early developmental stages (Capua & Eshed, 2017; Chandler et al., 2011; Durán-Medina et al., 2025; Ikeda et al., 2006; Nag et al., 2007). Moreover, *ESR2* expression appears to coincide with early stages of the establishment of new adventitious root primordia, and in basal root primordia. These observations beg the question of whether ESR2 may represent a more general founder cell or early organ primordia marker that also includes different types of roots. In accordance with the concept of extended stem cell niche in roots (Dubrovsky & Ivanov, 2021), cells in close proximity to the QC can also fulfill a role of stem cells. The fact that *ESR2* promoter activity was found in the primary root near the QC (**Fig. 8h**) raise a question whether *ESR2* plays a role in maintaining stem cell properties in roots. In conclusion, based on the experiments performed here on organogenetic capacities of different explants and regeneration conditions and intact loss-of-function and upon induction, ESR2 appears to impact organ formation in a context-dependent fashion. It promotes shoot regeneration, as expected, but it also inhibits or delays new root formation. It provides regenerative competence but should be downregulated or restricted at further stages for sustained root development. Moreover, *ESR2* is expressed at basal root primordia and when overexpressed delays rapid new basal root development after primary root removal. Furthermore, it promotes cell reprogramming for new root organogenesis in de-rooted hypocotyls. Finally, *ESR2* is also expressed at middle stages of LR formation in intact seedlings, specifically in the newly established QC in the developing LRP. Slow LRP formation in the *bol-cr2* mutant suggests that the *ESR2* expression in the QC is essential for LRP morphogenesis, In the same genetic background LR initiation is strongly inhibited underlying importance of ESR2 for LR formation and adding another function to this key organogenesis regulator.

The knowledge about genes that guide and participate in organogenesis programs supports the development of effective regeneration strategies that can open the door to genome edition of a broader range of plant species. Moreover, it provides a strong basis for strategies oriented to optimize organ-related and general plant architecture traits.

## SUPPLEMENTARY MATERIAL

**Supplementary Figure 1.** Effects of ESR2 in the root explant one-step regeneration assay in SIM.

**Supplementary Figure 2.** Effects of ESR2 in the root explant one-step regeneration assays in RIM.

**Supplementary Figure 3.** Effects of ESR2 in the root explant two-step regeneration assay in RIM.

**Supplementary Figure 4.** Effects of ESR2 in hypocotyl explant one-step regeneration assays.

**Supplementary Figure 5.** ESR2 expression during adventitious root formation from leaf explants.

**Supplementary Figure 6.** Basal and adventitious root emergence in de-rooted aerial explants.

**Supplementary Figure 7.** Summary of ESR2 effects in regeneration and organogenesis from diverse explants and in intact plants.

## ACKNOWLEDGEMENTS

We thank the anonymous reviewers for their insights to improve the manuscript. We also thank Ana Cecilia Ascencio, Brenda Marlene Valdivia Duarte, and Marian Wendolin Garcia Jimenez for technical assistance for phenotypic measurements and help with the experiments, and the Laboratory of Learning and Research in Computational Biology (Laicbio) at Cinvestav Irapuato Unit for access to their resources.

## AUTHOR CONTRIBUTIONS

Conceptualization: H.G.-L., J.G.D., N.M.-M; Experimental design: H.G.-L., M.I.C.E.-A., C.M.M.-P., H.M.J.-J., J.G.D., N.M.-M.; Experimental performance: H.G.-L., B.E.R.C, C.M.M.-P., M.I.C.E.-A, H.M.J.-J., J.G.D.; Formal analysis: H.G.-L., J.G.D.; Writing-initial draft preparation: H.G.-L.,; Figure preparation: H.G.-L., B.E.R.C.; Writing-review and editing: H.G.-L., N.M.-M.; Final manuscript version: J.G.D., H.G.-L., N.M-M.; Funding acquisition and resources: N.M.-M. and J.G.D.; Supervision: N.M.-M. All authors read and approved the final manuscript.

## DECLARATIONS

### Ethical Approval

This study did not involve any human participants or animals, and no ethical approval was required.

Herenia Guerrero-Largo

Ma. Isabel Cristina Elizarraraz-Anaya

Cecilia Monserrat Martínez-Pérez

Beatriz Ruiz-Cortés

Henri Manuel Jimenez-Jimenez

Joseph G. Dubrovsky

Nayelli Marsch-Martínez

