## Supplementary figures for "The transcription factor ESR2/DRNL/BOL differentially regulates *de novo* organogenesis, regeneration, and lateral root development in *Arabidopsis thaliana*"

Guerrero-Largo et al.

**Supplementary Figure 1.**

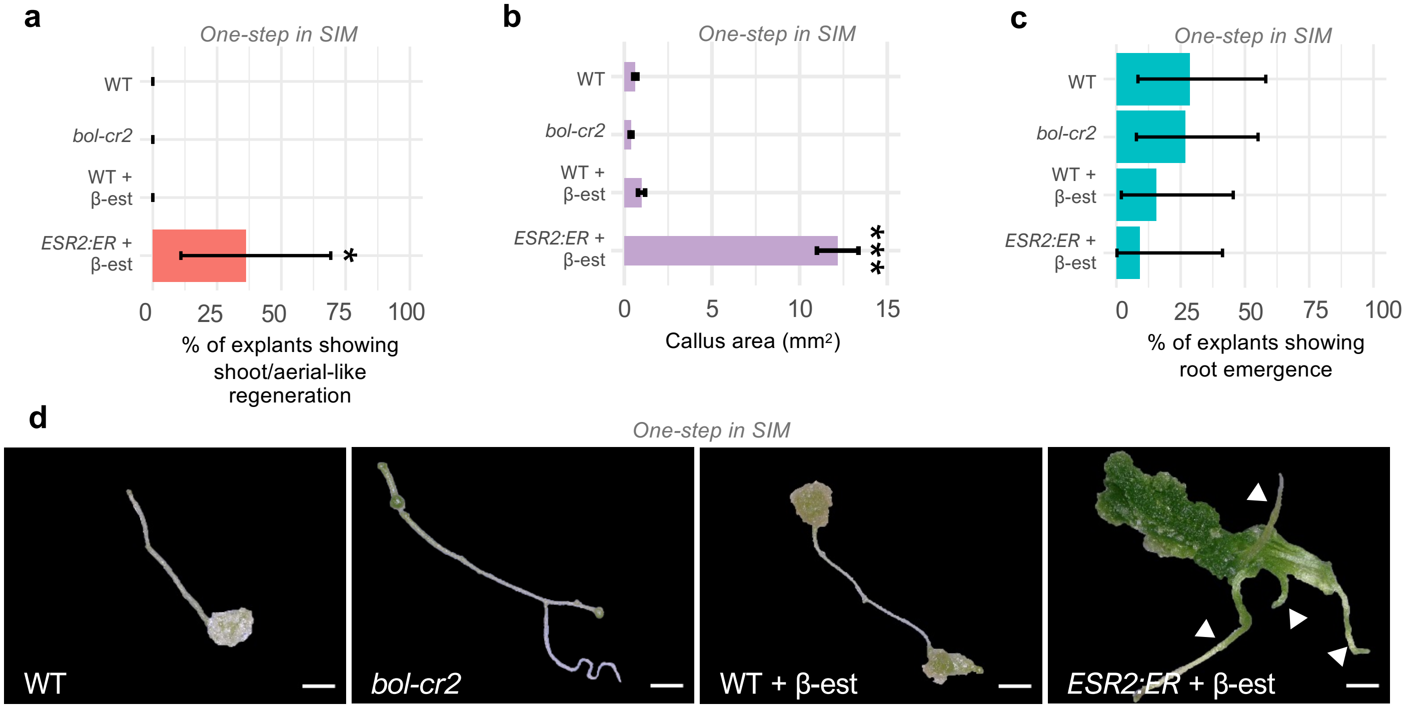


**Supplementary Figure 1. Effects of ESR2 in the root explant one-step regeneration assay in SIM. a** Frequency of shoot/aerial-like regeneration quantified 20 days after transference to SIM. Error bars indicate 95% confidence intervals calculated using an exact binomial method. Statistical significance was assessed using a test for binomial proportions (* $p<0.05$). **b** Callus area. Bars represent mean. Error bars represent SE. Statistical significance was determined by *t*-test (*** $p<0.001$). **c** Frequency of explants showing root emergence. Error bars indicate 95% confidence intervals calculated using an exact binomial method. **d** Representative root explants of each genotype. White arrowheads indicate stem-like structures. Scale bars = 1 mm. β-est: β-estradiol. **a-c** $n$ = 12-17 seedlings per line, two independent experiments.

**Supplementary Figure 2.**

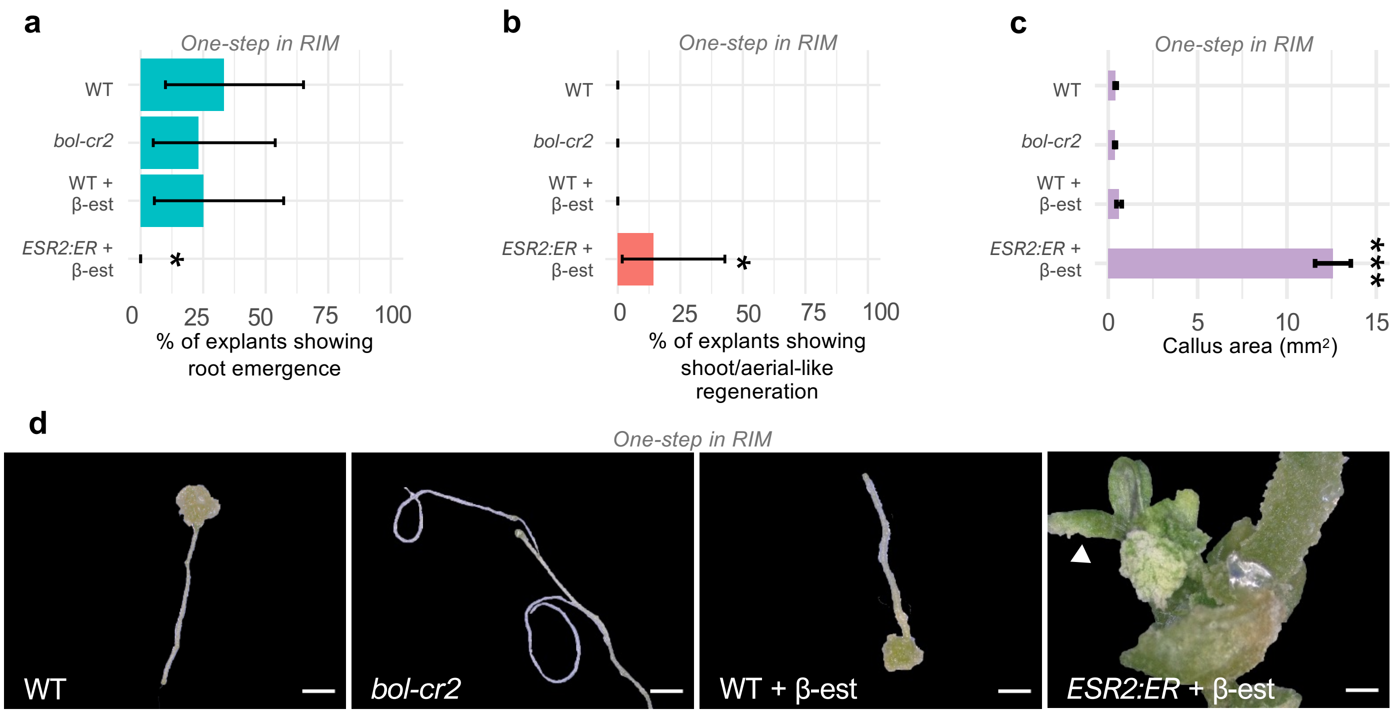


**Supplementary Figure 2. Effects of ESR2 in the root explant one-step regeneration assays in RIM. a-b** Frequency of root explants showing shoot/aerial-like (a) and root emergence (b) quantified 20 days after transference to Root Inducing Medium (RIM). Error bars indicate 95% confidence intervals calculated using an exact binomial method. Statistical significance was assessed using a test for binomial proportions (* $p$<0.05). **c** Callus area. Bars represent mean. Error bars represent SE. Statistical significance was determined by $t$-test (*** $p$<0.001). **d** Representative root explants of each genotype. Scale bars = 1 mm. β-est: β-estradiol. ). **a-c** n = 12-17 seedlings per line, two independent experiments.

**Supplementary Figure 3.**

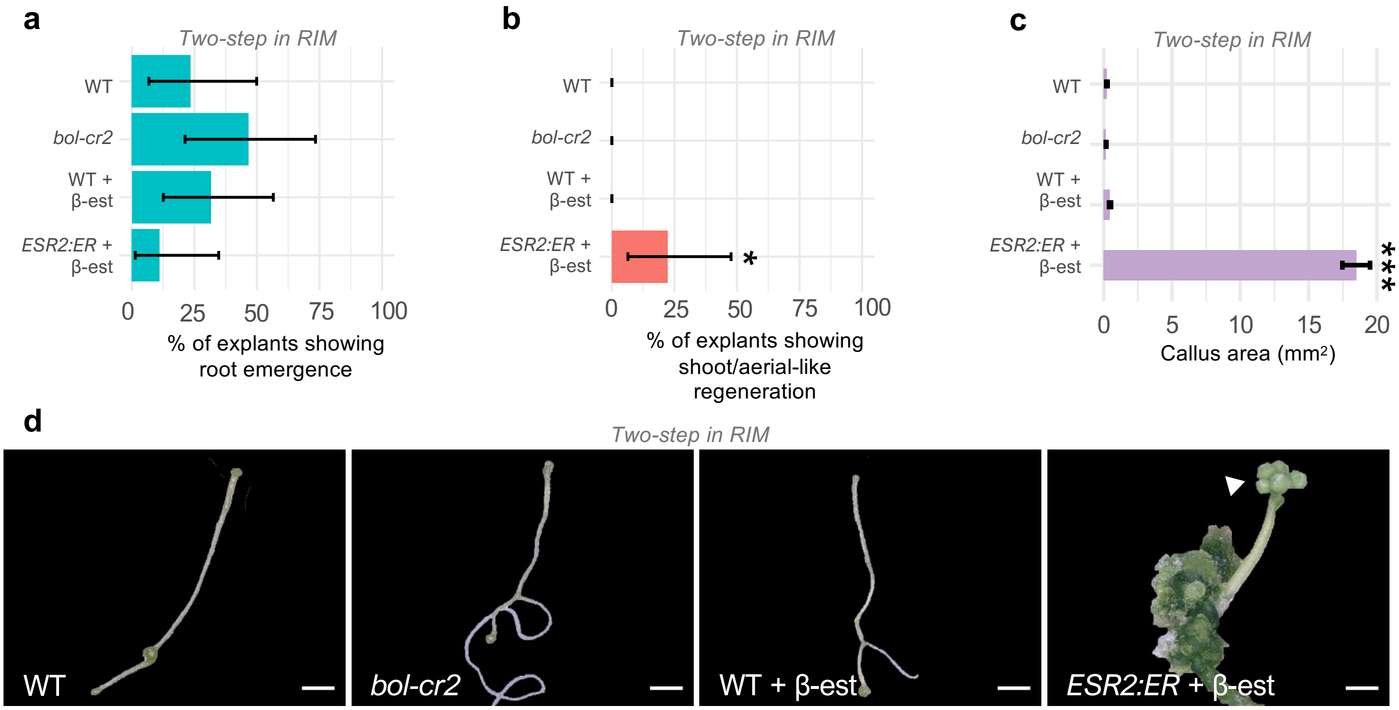


**Supplementary Figure 3. Effects of ESR2 in the root explant two-step regeneration assay in RIM. a-b** Frequency of root explants showing shoot/aerial-like (a) and root emergence (b) quantified 20 days after transference to Root Inducing Medium (RIM). Error bars indicate 95% confidence intervals calculated using an exact binomial method. Statistical significance was assessed using a test for binomial proportions (* $p$<0.05). **c** Callus area. Bars represent mean. Error bars represent SE. Statistical significance was determined by $t$-test (*** $p$<0.001). **d** Representative root explants of each genotype. Scale bars = 1 mm. β-est: β-estradiol. ). n = 12-17 seedlings per line, two independent experiments.

**Supplementary Figure 4.**

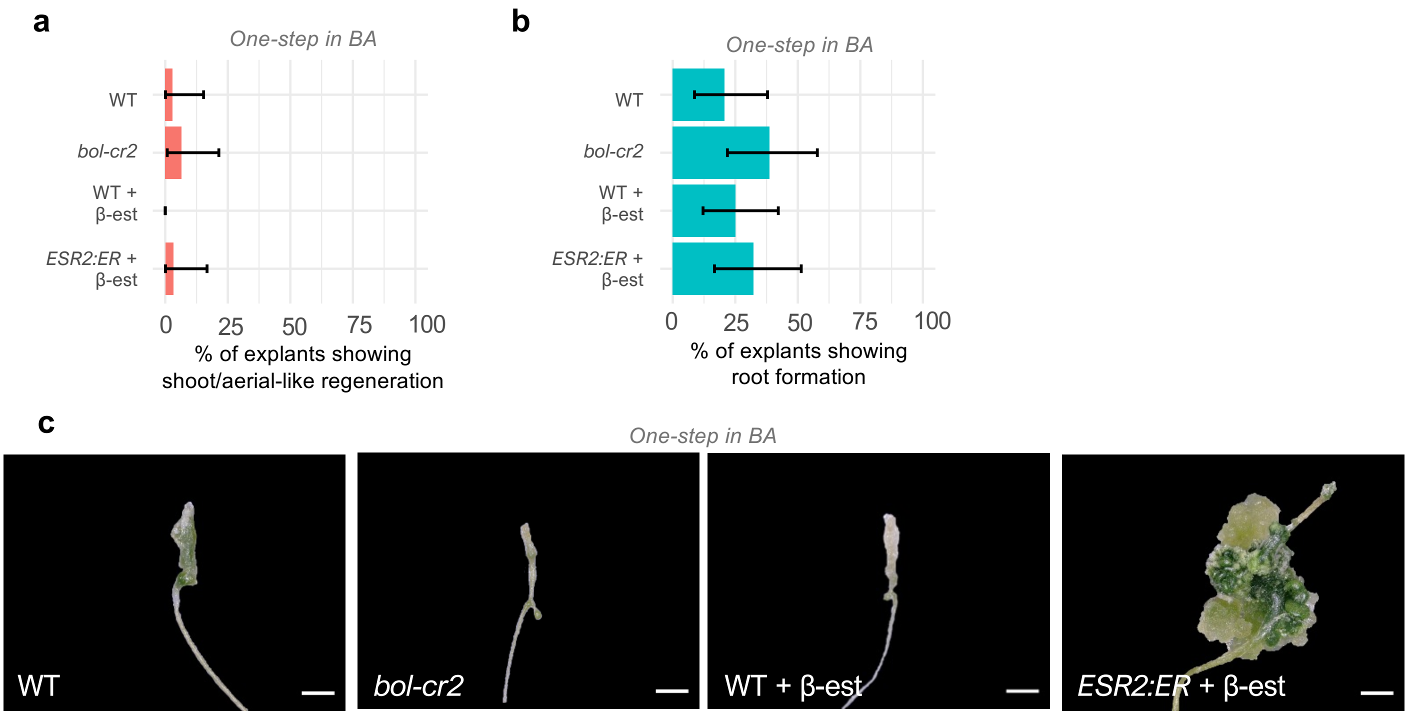


**Supplementary Figure 4. Effects of ESR2 in hypocotyl explant one-step regeneration assays. a-b** Frequency of explants showing shoot/aerial-like (a) and root formation (b) quantified 20 days after transfer to medium supplemented with Benzylaminopurine (BA). Error bars indicate 95% confidence intervals calculated using an exact binomial method. Statistical significance was assessed using a test for binomial proportions ($p$>0.05). n = 31-36 seedlings per line, two independent experiments. **c** Representative root explants without shoot regeneration of each genotype. Scale bars = 1 mm. β-est: β-estradiol.

**Supplementary Figure 5.**

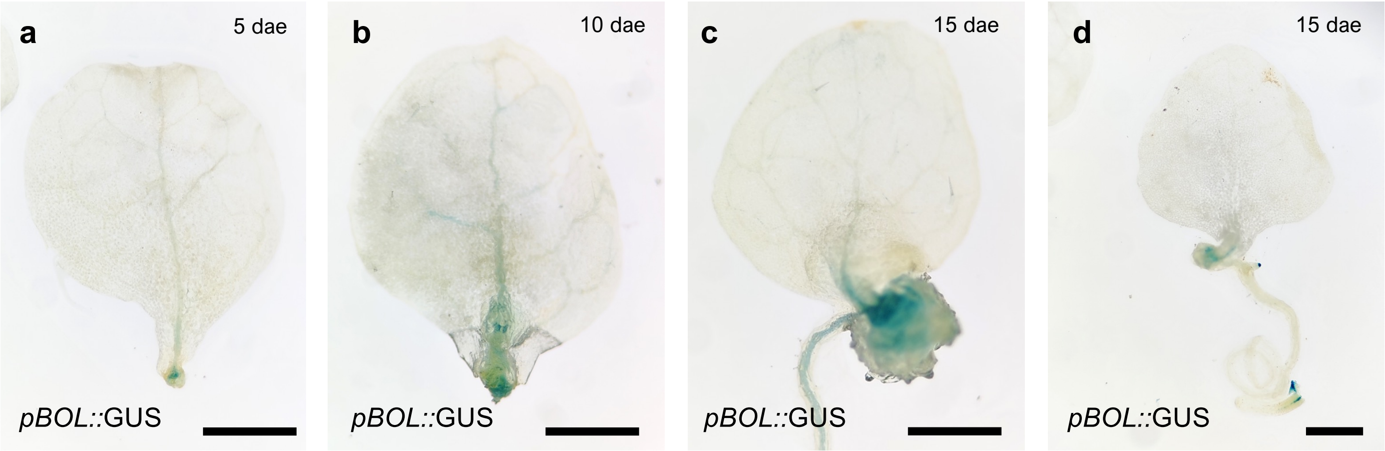


**Supplementary Figure 5. *ESR2* expression during adventitious root formation from leaf explants.** *ESR2* expression at five **(a)**, ten **(b)**, and fifteen **(c-d)** days after excision (dae) and transference onto B5 Gamborg's medium. Scale bars = 1 mm.

**Supplementary Figure 6.**

**a**

| | **Total roots** | | | **Basal roots** | | | **Adventitious roots** | | |
|---|---|---|---|---|---|---|---|---|---|
| | Mean | SD | *P* value | Mean | SD | *P* value | Mean | SD | *P* value |
| WT | 1.9 | 0.6 | | 1.9 | 0.6 | | 0 | 0 | |
| *bol-cr2* | 2.5 | 0.7 | *** | 2.5 | 0.7 | *** | 0 | 0 | |
| WT+β-est | 1.8 | 0.6 | | 1.8 | 0.6 | | 0 | 0 | |
| *ESR2:ER* + β-est | 3.0 | 0.9 | *** | 1.6 | 0.9 | | 1.2 | 0.9 | *** |

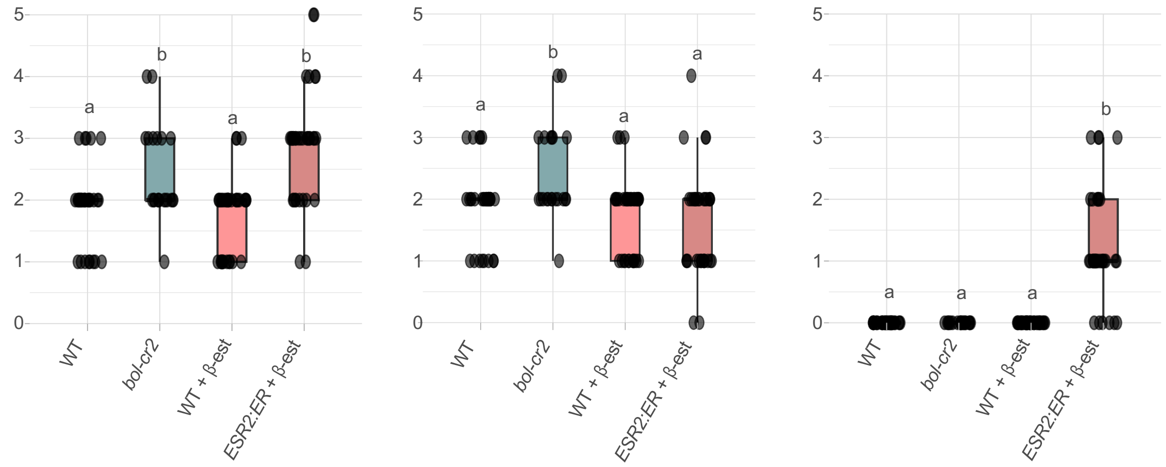


**b**

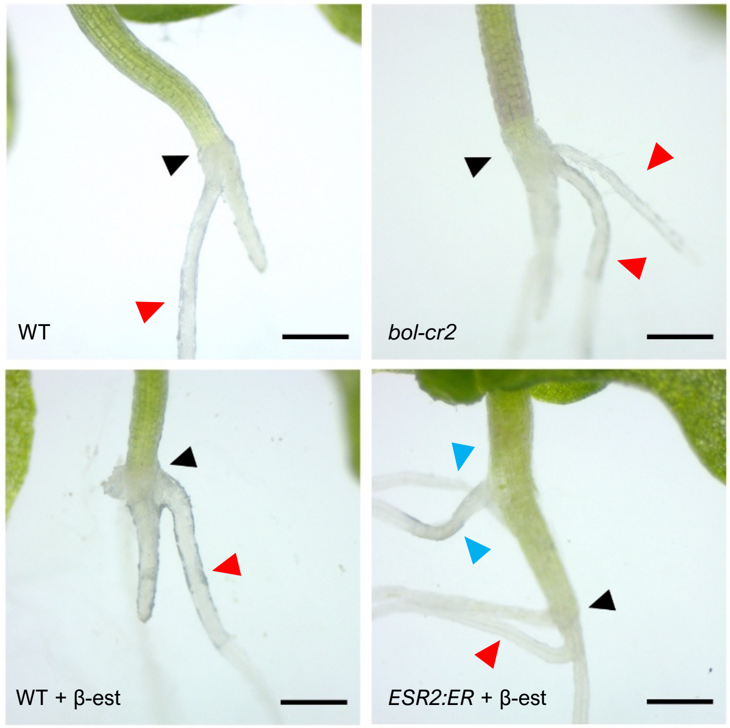


**c**

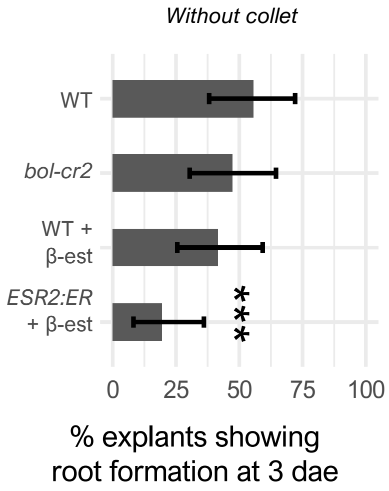

**Supplementary Figure 6. Basal and adventitious root emergence in de-rooted aerial explants.** **a** Table and graphs summarize statistic values of total, basal and adventitious root number across genotypes in de-rooted explants with collet at 6 dae (*** $p<0.001$, ANOVA/Tukey test). n =  seedlings per line, two independent experiments. **b** Representative images of de-rooted aerial explants with collet region 6 dae transferred onto B5 Gamborg's medium. Note that roots that appear as a continuation of the hypocotyl axis are roots that regenerated from a collet region. Black arrowhead = collet; red = basal root; blue = adventitious root. β-est: β-estradiol. Scale bars = 500 µm. **c** Frequency of explants without collet showing root formation 3 days after excision (dae)**.** Error bars indicate 95% confidence intervals calculated using an exact binomial method (*** $p<0.001$, test for binomial proportions). n = 33-36 seedlings per line, two independent experiments.

**Supplementary Figure 7.**

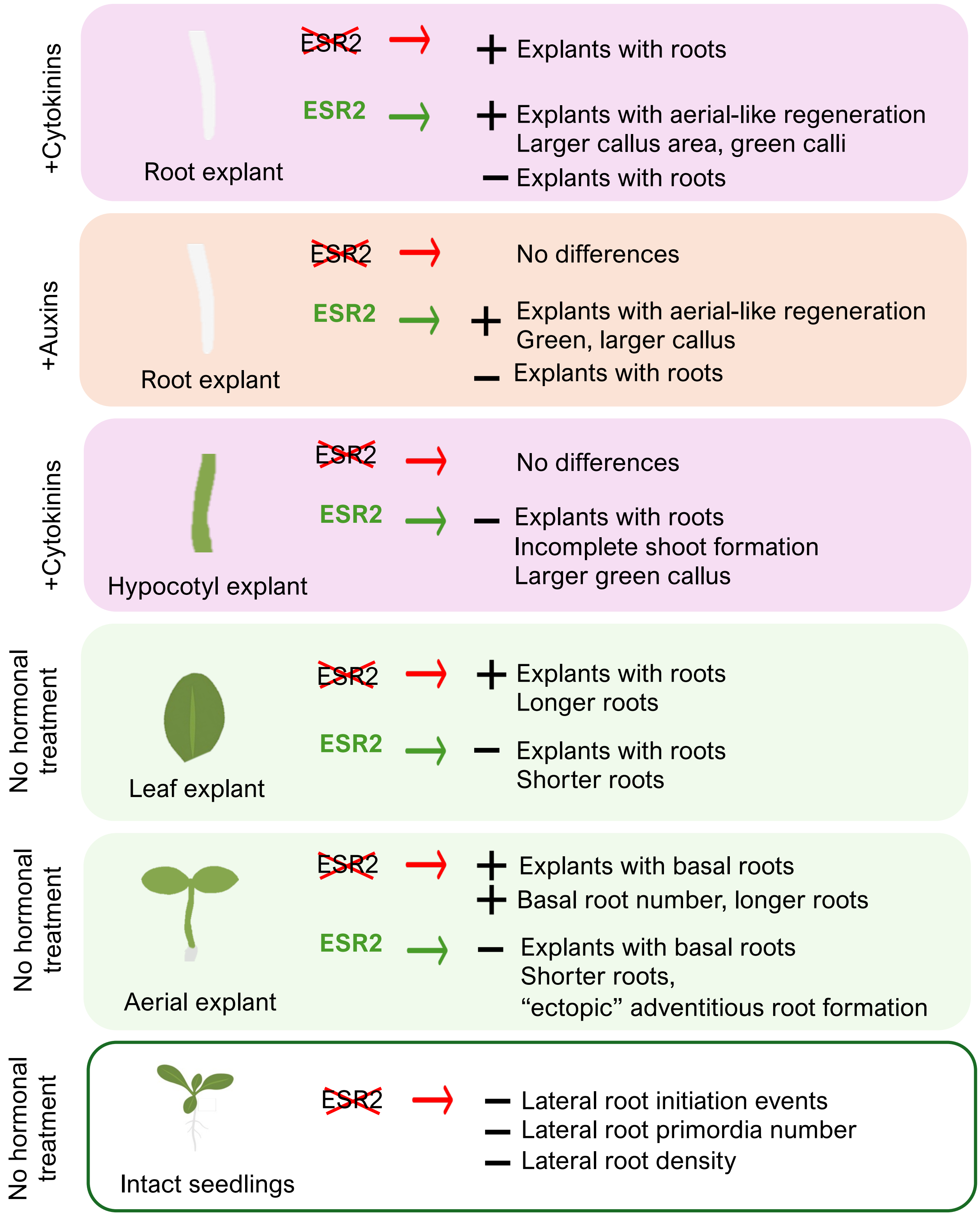

**Supplementary Figure 7. Summary of ESR2 effects in regeneration and organogenesis from diverse explants and in intact plants.** The blocks summarize the effect of the loss of function or induced ESR2 activity backgrounds compared to wild type in different contexts. The last block indicates the effects of the loss of ESR2 function in lateral root formation in intact plants. The red ”x” indicates loss of function and the bold green letters indicate overexpression.
